# Chromatin accessibility landscapes activated by cell surface and intracellular immune receptors

**DOI:** 10.1101/2020.06.17.157040

**Authors:** Pingtao Ding, Toshiyuki Sakai, Ram Krishna Shrestha, Nicolas Manosalva Perez, Wenbin Guo, Bruno Pok Man Ngou, Shengbo He, Chang Liu, Xiaoqi Feng, Runxuan Zhang, Klaas Vandepoele, Dan MacLean, Jonathan DG Jones

## Abstract

Activation of cell Surface and Intracellular Receptor-Mediated Immunity (SRMI and IRMI) results in rapid transcriptional reprogramming that underpins disease resistance. However, the mechanisms by which SRMI and IRMI lead to transcriptional changes are not clear. Here, we combine RNA-seq and ATAC-seq to define changes in gene expression and chromatin accessibility; both SRMI and IRMI increase chromatin accessibility at induced defense genes. Analysis of ATAC-seq and RNA-seq data combined with publicly available information on transcription factor DNA-binding motifs enabled comparison of individual gene regulatory networks activated by SRMI and IRMI, and by both. These results and analyses reveal overlapping and conserved transcriptional regulatory mechanism between the two immune systems.

## Main

Plants use both cell-surface and intracellular receptors to detect pathogen-derived molecules and activate innate immunity^1^. Plant cell-surface immune receptors (pathogen recognition receptors, or PRRs) perceive relatively conserved pathogen-associated molecular patterns (PAMPs) or endogenous damage-associated molecular patterns (DAMPs) released from damaged or dying plant cells and activate Pattern- or DAMP-triggered immunity (PTI or DTI)^2,3^. Intracellular immune receptors in plants are usually nucleotide-binding, leucine-rich repeat (NLR)proteins. NLRs recognize, directly or indirectly, pathogen effectors secreted into plant cells and activate effector-triggered immunity (ETI). These innate immune systems involve distinct responses mediated by different subsets of molecular components^4^. Some cell-surface receptors, such as tomato Cf-4 and Cf-9 detect apoplastic effectors yet activate PTI-like responses^5^. The fundamental distinction is between processes initiated by PRRs or NLRs, so we refer to surface-receptor-mediated immunity (SRMI, pronounced as ‘surmi’) and intracellular-receptor-mediated immunity (IRMI, pronounced ‘ermi’) instead of PTI and ETI (Supplementary Fig. 1). In interactions between plants and microbial pathogens, SRMI will always precede IRMI, since effector delivery requires close host/microbe interaction.

We study the Arabidopsis RPS4/RRS1 NLR pair that detects bacterial effectors AvrRps4 and PopP2. Using a *Pseudomonas* strain that solely delivers one of these effectors, we defined early RPS4/RRS1-dependent transcriptional responses in Arabidopsis leaves^6,7^, and showed that 4 hours after infiltration, SRMI together with IRMI (‘SRMI+IRMI’) elevates the expression of defense-related genes more strongly compared to SRMI alone^6,8^. This early timepoint precedes accumulation of defense hormone salicylic acid (SA) and gene reprogramming in response to increased endogenous SA level^6,8^. This implies that IRMI-enhanced transcriptional regulation plays an essential role in conferring robust immune responses against pathogens^4^, but how IRMI activates defense genes remains unclear. To study IRMI-specific physiological changes, we generated an inducible IRMI system^9^.

Activation of SRMI, IRMI and ‘SRMI+IRMI’ lead to rapid transcriptional reprogramming^6,8,10^. Many transcriptional regulatory components are involved in orchestrating effective immunity in plants^11,12^, notably transcription factors (TFs)^13^, transcription co-repressors^14^, the Mediator complex^15^, histone-modifying enzymes^16^, and histone remodellers^17^. Little is known of how changes in transcription rates at defense genes are initiated, maintained and regulated upon the activation of either class of plant immune receptor.

Open or accessible chromatin regions (ACRs) at promoters and enhancers are associated with active gene expression in eukaryotes^18^. Applications of assays for transposase-accessible chromatin following by sequencing (ATAC-seq) in plants have revealed species-, tissue- and cell-type-specific chromatin signatures^19–23^ in recent studies, but chromatin accessibility changes associated with inducible responses, such as immune activation are less well characterized. We hypothesized that correlating immunity-specific transcriptomes with an atlas of open chromatin profiles could reveal novel *cis*-regulatory elements (CREs) and associated regulatory mechanisms. We therefore performed a set of comparative analyses with ATAC-seq and RNA-seq data generated during SRMI, ‘SRMI+IRMI’ and IRMI. This study provides a direct link between changes in chromatin accessibility and associated gene expression and new insights into the dynamics of chromatin accessibility landscapes and gene regulatory networks during plant immune activation.

### ATAC-seq in Arabidopsis reveals tissue-specific chromatin accessibility

ATAC-seq was first used to capture open chromatin regions in human cell lines and rapidly adapted to other eukaryotic systems including plants^19,24^. To study the dynamic chromatin features during plant immune activation, we established a protocol to prepare fresh nuclei isolated from adult rosette leaves of Arabidopsis (*Arabidopsis thaliana*) Columbia-0 (Col-0) ecotype using fluorescence-activated nuclei sorting (FANS) (Supplementary Fig. 2a). A similar approach was reported previously^19^. To generate FANS-ATAC-seq libraries from multiple samples that are (i) compatible with the Illumina next-generation sequencing (NGS) sequencing platforms and (ii) can be multiplexed we designed and synthesized barcoded primers with 9-nucleotide (nt) unique indices for dual index and paired-end sequencing (Supplementary Fig. 2b-d; Supplementary Table 1). In a trial run we used 10k, 20k, 50k and 80k sorted nuclei as ATAC input with a fixed amount of ‘tagmentation’ reaction, to obtain an optimal ratio between the input nuclei (DNA) and Tn5 transposase (Supplementary Fig. 3a). Purified naked Arabidopsis genomic DNA was tagmented in three replicates and sequenced as controls for ATAC-seq normalization (Supplementary Fig. 2d). In this trial FANS-ATAC-seq run, we observed reproducible accessible chromatin features captured in two biological replicates with different levels of input (Supplementary Fig. 3b-e).

To test if this ATAC-seq method is sensitive enough to detect tissue-specific chromatin accessible features, we additionally performed FANS-ATAC-seq with sperm nuclei (SN) and vegetative nuclei (VN), the male germ unit derived from Arabidopsis pollen grain. We found that ACRs enriched at the *SYSTEMIC ACQUIRED RESISTANCE DEFICIENT 1* (*SARD1*) defense gene locus are only observed in somatic but not germline cells (Supplementary Fig. 4a). *SARD1* encodes a TF involved in plant immunity^13,25,26^. We inspected another well-known *Resistance (R)-*gene cluster on Arabidopsis chromosome 4 which harbors a group of N-terminal Toll/interleukin-1 receptor/resistance protein (TIR) domain-containing NLRs. Similar to *SARD1*, promoters of *RECOGNITION OF PERONOSPORA PARASITICA 4* (*RPP4*) or *CHILLING SENSITIVE 2* (*CHS2*), *SUPPRESSOR OF NPR1-1, CONSTITUTIVE 1* (*SNC1*), *SIDEKICK SNC1 1* (*SIKIC2*) and *RESISTANCE TO LEPTOSPHAERIA MACULANS 3* (*RLM3*) show enriched ACRs in leaf nuclei ATAC-seq data compared to the other four NLRs in the same gene cluster, but not in SN or VN ATAC-seq data (Supplementary Fig. 4b). This is consistent with the observation that expression levels of *RPP4/CHS2, SNC1, SIKIC2* and *RLM3* in Arabidopsis adult leaves are much higher than the other four NLRs in the same gene cluster^27^. Expression of *RPP4/CHS2, SNC1, SIKIC2* and *RLM3* in Arabidopsis leaves contributes to resistance against multiple pathogens^28–32^. In addition, trimethylation of the 4^th^ lysine of the histone H3 (H3K4me3s), histone marks that are often associated with actively transcribed genes, is enriched in *RPP4/CHS2* and *SNC1* promoters in Arabidopsis^33^, supporting our ATAC-seq results (Supplementary Fig. 4b). Overall, ACRs enriched in immunity-related genes are specific to somatic but not to germline cells.

### ATAC-seq to study Arabidopsis inducible innate immunity

We applied the FANS-ATAC-seq method to study changes in chromatin accessibility associated with gene expression induced by innate immunity. In Arabidopsis Col-0, two paired NLR proteins RPS4/RRS1 and RPS4B/RRS1B serve as intracellular NLR receptors activating IRMI upon recognition of AvrRps4, an effector derived from *Pseudomonas* (*P*.) *syringae* pv. *pisi*, a causal agent of bacterial blight in pea (*Pisum sativum*)^34^. We use a non-pathogenic strain of *P. fluorescens* Pf0-1 engineered with the type III secretion system (T3SS) from *P. syringae* (‘Effector-to-Host Analyzer’ or EtHAn) tool to deliver wild-type AvrRps4 (Pf0-1:AvrRps4^WT^) or its mutant (Pf0-1:AvrRps4^mut^) into Col-0 leaf cells^35,36^. AvrRps4^mut^ (KRVY135-138AAAA) is unable to activate IRMI mediated by RPS4/RRS1 and RPS4B/RRS1B^34^. Infiltration of Pf0-1:AvrRps4^mut^ activates SRMI, and Pf0-1:AvrRps4^WT^ activates ‘SRMI+IRMI’ (Supplementary Fig. 5a), as in previous reports^4,8^. We took samples at 4 hours post-infiltration (hpi) for ATAC-seq to monitor early changes during immune activation (Supplementary Fig. 5a)^6,8^. We obtained similar genome-wide ATAC-seq peak coverage patterns with different treatments (Supplementary Fig. 5b).

ATAC-seq peaks in all biological replicates under different conditions were enriched within 2 kilobases (kb) upstream of the transcription start site (TSS) and within 1 kb downstream of the transcript termination site (TTS) (Fig. 1a,b). The distribution of ACRs relative to genomic features was highly similar between all ATAC-seq data sets (Supplementary Fig. 5c-f; Supplementary Table 2). Over 77% of ACRs are mapped to the putative gene promoters (pACRs; within 2 kb upstream of a gene) (Supplementary Fig. 5c-f), consistent with previously reported ATAC-seq data sets^19,37^. ∼8% of ACRs mapped to distal intergenic genome loci (dACRs) (Supplementary Fig. 5c-f), slightly higher than 5.9% reported recently^23^. In addition, compared to 16,296 ACRs observed in total using a similar FANS-ATAC-seq approach in a recent report^23^, we obtained a range of 24,901 to 27,285 total ACRs (Fig. 1c; Supplementary Table 3), also slightly more than the 23,288 total reported elsewhere applying ATAC-seq with INTACT (isolation of nuclei tagged in specific cell types) purified nuclei^20^. Comparing ACRs enriched in all conditions, we found 10,658 (∼ 40% of total ACRs) are shared (Fig. 1c). The remaining 60% unshared ACRs may point to specific regulatory signals under each condition.

**Figure 1.**
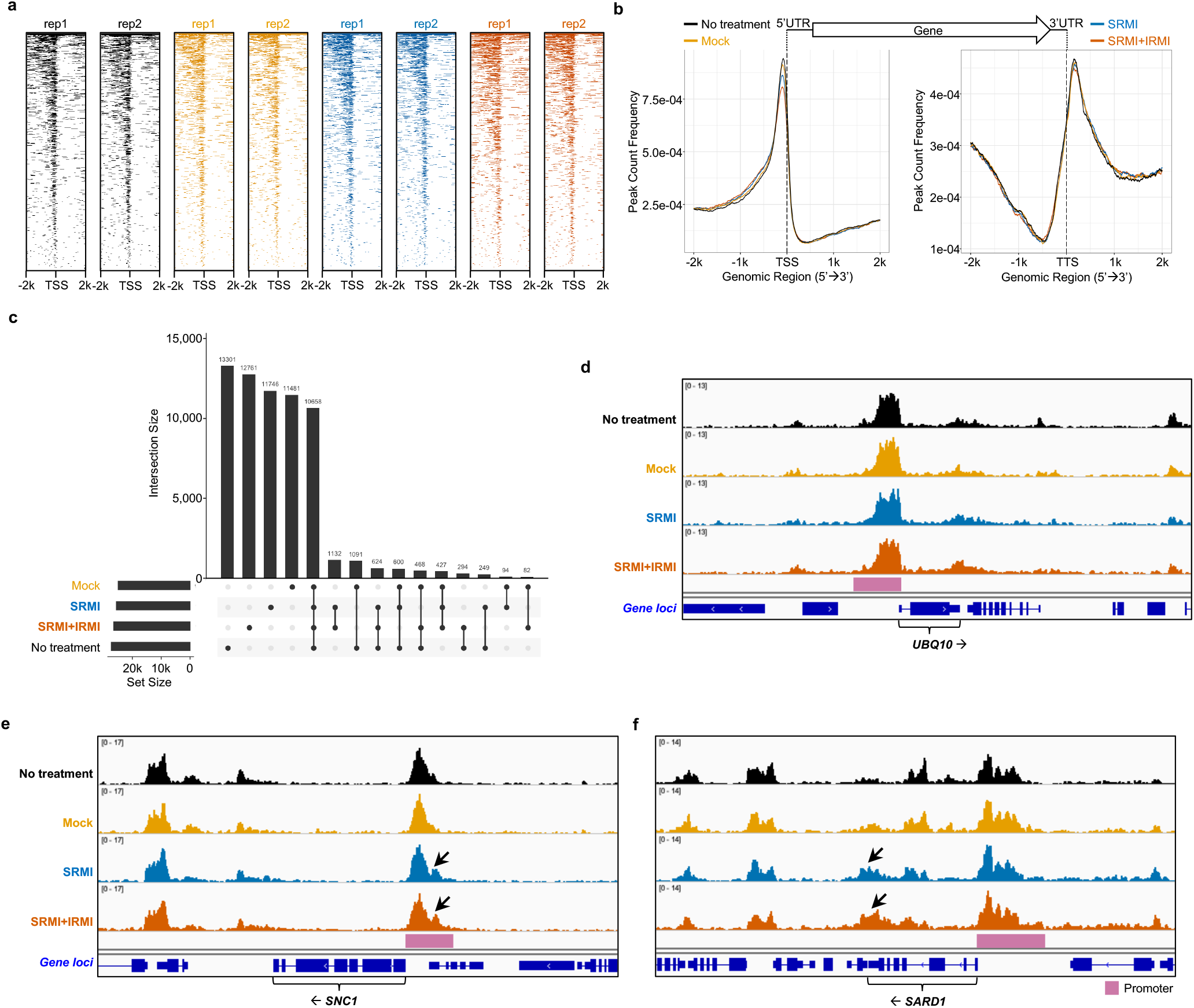
Interrogation of chromatin landscapes activated by SRMI and ‘SRMI+IRMI’. **(a)** Heatmaps showing the distribution of accessible regions around the TSS identified by FANS-ATAC-seq in two biological replicates (rep) under four different conditions. Accessible regions are mapped to 2, 000 bp upstream (−2k) or downstream (2k) of TSS as the center. ‘No treatment’, Mock, SRMI and ‘SRMI+IRMI’ are indicated in black, orange, blue and vermillion, respectively (same color codes apply to the same set of treatments in the rest of this study). **(b)** Distribution of accessible regions around the TSS (left panel) and TTS (right panel) identified from FANS-ATAC-seq with the mean peak counts from two biological replicates indicated in (a). The center of accessible regions was used to produce the distribution plots. **(c)** An UpSet plot showing the relationships of accessible regions enriched under four different conditions indicated in (a) and (b). ‘Intersection Size’ indicates either condition specific accessible regions or shared accessible regions under different combinations of condition comparisons. ‘Set Size’ indicates the aggregates or total number of accessible regions found under each condition. **(d-f)** Genome browser views of ATAC-seq-indicated chromatin accessibility changes occurring near selected gene loci under different conditions. Gene symbols are labeled for the corresponding gene loci, (d) *UBQ10*, (e) *SNC1* and (f) *SARD1*. The protein coding strand is indicated in black arrow placed next to the gene symbol. The reddish-purple bars next to each gene loci indicate their putative promoter regions. **Related supplementary information:** Supplementary Fig. 5 Supplementary Tables 2 and 3

Among those shared ACRs, pACRs enriched at house-keeping gene loci, such as the *UBQ10* (*POLYUBIQUITIN 10*), show similar patterns in all conditions (Fig. 1d), consistent with the presumed constitutive expression. pACRs enriched at *SNC1* and *SARD1* are similar to those observed in our trial run (Supplementary Fig. 4a,b) and the major peaks of pACRs at these two gene loci under different conditions are similar. We observe additional small pACRs at *SNC1* and increased ACRs at the 3’UTR of *SARD1* upon SRMI and ‘SRMI+IRMI’ treatment (Fig. 1e,f). Those observations are positively correlated with previous reports that expression of *SARD1* and *SNC1* are upregulated by immune activation^8,33^.

### Positive correlation of increased ACRs and expression of defense genes during SRMI and ‘SRMI+IRMI’

We reported that some defense genes are induced by both SRMI and ‘SRMI+IRMI’ by profiling expressions of selected genes^8^. In this study, we performed genome-wide RNA-seq to study genes induced by SRMI and ‘SRMI+IRMI’ more extensively (Supplementary Fig. 6a,b). There are 4665 and 5004 upregulated genes during SRMI and ‘SRMI+IRMI’ compared to the ‘No treatment’ control, respectively. Among these, 4494 genes are shared by SRMI and ‘SRMI+IRMI’ (Supplementary Fig. 6c, Supplementary Tables 4). Similarly, there are 5433 downregulated genes shared by SRMI and ‘SRMI+IRMI’ (Supplementary Fig. 6c). This greatly expands the shared list of genes showing similar regulatory patterns between SRMI and ‘SRMI+IRMI’ compared to our previous report (Supplementary Fig. 6c,d; Supplementary Tables 5)^8^. Upregulated genes shared by SRMI and ‘SRMI+IRMI’ are mostly enriched in gene ontology as defense-related genes or genes in response to stress (Supplementary Fig. 6e,g), whereas downregulated genes shared by both immune activation conditions are enriched with respect to genes involved in photosynthesis (Supplementary Fig. 6f,h). These results indicate a transcriptional reprogramming from photosynthesis to defense activation in Arabidopsis leaves upon activation of both immune conditions. In addition, we identified 3005 genes that are more strongly induced by ‘SRMI+IRMI’ compared to SRMI alone, and they distribute in Clusters 5, 7 and 9 based on their co-expression pattern (Supplementary Fig. 6d, i-k).

We hypothesized that rapid elevation of gene expression would be correlated with increased chromatin accessibility at these gene loci during immune activation, as active transcription usually requires increased access of DNA-binding proteins such as TFs and transcriptional machineries^38^. To test this, we displayed ATAC-seq and corresponding RNA-seq data for well-known defense gene loci (Fig. 2). These were *ISOCHORISMATE SYNTHASE 1* (*ICS1*), *ENHANCED DISEASE SUSCEPTIBILITY 5* (*EDS5*) and *AVRPPHB SUSCEPTIBLE 3* (*PBS3*), genes involved in SA biosynthesis^39–42^; and *AGD2-LIKE DEFENSE RESPONSE PROTEIN 1* (*ALD1*), *SAR DEFICIENT 4* (*SARD4*) and *FLAVIN-DEPENDENT MONOOXYGENASE 1* (*FMO1*), genes involved in synthesizing pipecolic acid (Pip) and its derivatives, that contribute to systemic acquired resistance (SAR) and defense priming^43–46^. We observed strongly increased ACRs at the promoters of all six selected genes in SRMI and ‘SRMI+IRMI’ compared to ‘No treatment’ or Mock treatments (Fig. 2a-f) and increased transcripts of those genes (Fig. 2g-i). This indicates a positive correlation between rapid transcriptional upregulation of selected defense genes and increased ACRs near the corresponding gene loci during activation of SRMI and ‘SRMI+IRMI’.

**Figure 2.**
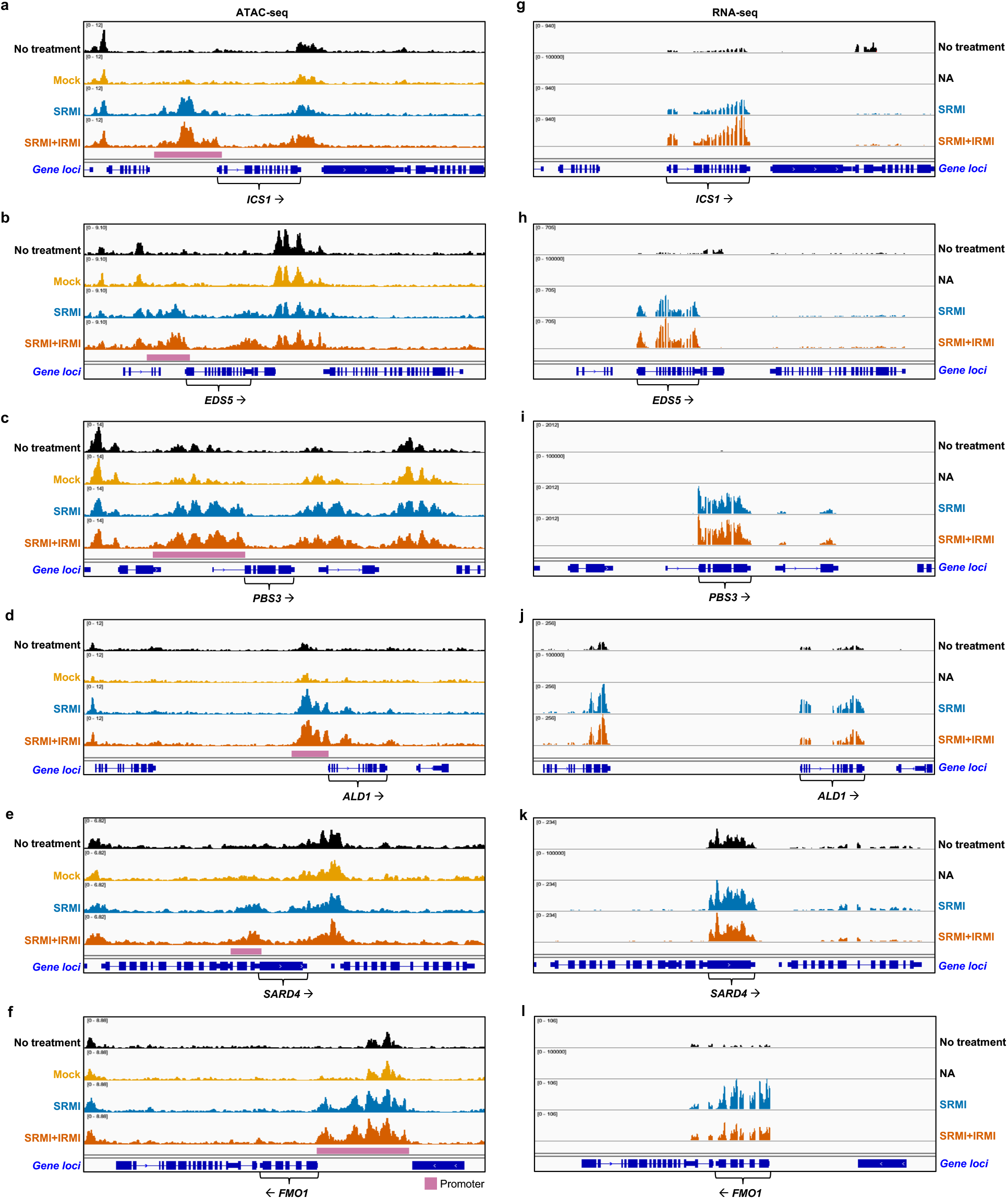
Integration of ATAC-seq with RNA-seq for genes activated by SRMI and ‘SRMI+IRMI’. **(a-f)** and **(g-i)** Genome browser views of ATAC-seq and RNA-seq on indicated gene loci under different conditions. **(a)** and **(g)** are date for *ICS1* locus. **(b)** and **(h)** are date for *EDS5* locus. **(c)** and **(i)** are date for *PBS3* locus. **(d)** and **(j)** are date for *ALD1* locus. **(e)** and **(k)** are date for *SARD4* locus. **(f)** and **(l)** are date for *FMO1* locus. *ICS1, EDS5* and *PBS3* are genes encoding enzymes that are required for the biosynthesis of a defense hormone, salicylic acid (SA) in the isochorismate pathway. *ALD1, SARD4* and *FMO1* are enzyme encoding genes involved in the biosynthesis of a secondary metabolite, N-hydroxy pipecolic acid (NHP) that are required to initiate systemic acquired resistance in plants. ATAC-seq data contain the same four conditions as shown in Figure 1, whereas the data point of Mock treatment is absent in RNA-seq data (NA). ATAC-seq and RNA-seq data points with the same corresponding labels indicate they were collected under exactly the same conditions (see more details in Methods). **Related supplementary information:** Supplementary Fig. 6 Supplementary Tables 4 and 5

### Genome-wide assessment of gene regulatory changes during IRMI

Activation of IRMI requires effector delivery from a pathogen, so will usually be preceded by SRMI (except perhaps for some recognized viruses). Previous studies on ETI (aka IRMI) usually involve effector delivery from *Pseudomonas sp*. or *Agrobacterium* transient expression, and thus are studies on ‘SRMI+IRMI’. We reported previously an inducible IRMI system by expressing AvrRps4^WT^ (SETI^WT^) in which AvrRps4^WT^ is only expressed upon β-estradiol (E2) induced nuclear binding of XVE to the LexA operon (E2:AvrRps4^WT^)^9^. This system enables investigation of IRMI-specific physiological changes^4^.

IRMI induced in SETI^WT^ displays similar transcriptional dynamics to that induced by Pf0-1 EtHAn^9^. We focused on IRMI-specific transcriptional activation; all RNA-seq samples were collected at a relatively early time point of the activation (4 hpi of E2) (Supplementary Fig. 7a)^4,9^. To obtain the list of differentially expressed genes (DEGs) during IRMI, we compared gene expression profiles in E2-treated SETI^WT^ at 4 hpi (IRMI) to those in E2-treated SETI^WT^ at 0 hpi (Control_1) or to those in E2-treated SETI^mut^ at 4 hpi (IRMI_mut) (Supplementary Fig. 7a-d; Supplementary Tables 6 and 7). The comparisons of ‘IRMI vs Control_1’ and ‘IRMI vs IRMI_mut’ share mostly the same genes in both up- and down-regulation groups (1584 shared upregulated and 1869 shared downregulated genes, respectively) (Supplementary Fig. 7b-d). The number of DEGs in ‘IRMI vs Control_1’ is much more than that in ‘IRMI vs IRMI_mut’ (Supplementary Fig. 7b,d). The majority of up- and down-regulated DEGs in ‘IRMI vs IRMI_mut’ were shared by ‘IRMI vs Control_1’ (Supplementary Fig. 7c; Supplementary Tables 6). From the gene ontology (GO) enrichment analysis of DEGs in those comparisons, we found the GO term of ‘response to wounding’ is enriched in DEGs of ‘IRMI vs Control_1’ but not in that of ‘IRMI vs IRMI_mut’ (Supplementary Fig. 7e-h). This indicates that both IRMI and IRMI_mut activate genes that are induced by mechanical wounding via the infiltration process at 4 hpi. Thus, comparing IRMI to IRMI_mut in ‘IRMI vs IRMI_mut’ eliminates wounding-induced genes, and reduces background. From these DEGs in ‘IRMI vs IRMI_mut’, we found genes mostly enriched in GO terms of ‘response to chitin’, ‘protein phosphorylation’ and ‘defense response’, (Supplementary Fig. 7i-k)^4^.

To study the changes in accessible chromatin on the loci of DEGs, we performed FANS-ATAC-seq (Supplementary Fig. 8a). Instead of using E2 treatment at 0 hpi, we use mock-treated sample at 4 hpi as a negative control, imitating the effects from wounding (Supplementary Fig. 8a). We observed consistent genomic distribution patterns of ACRs in all samples (Supplementary Fig. 8b-j; Supplementary Tables 8). To demonstrate ACRs that are specifically induced by IRMI compared to all control conditions, we checked the *ICS1* locus in comparison to the house-keeping gene *UBQ10* locus (Fig. 3a,b). We found only IRMI induces DARs at *ICS1* promoter and 3’UTR, but not in the ‘No treatment_1’ control (Fig. 3a). In contrast significant DARs are induced at *UBQ10* promoter and proximal region among all treatments (Fig. 3b). This is consistent with stable expression of *UBQ10* under all conditions.

**Figure 3.**
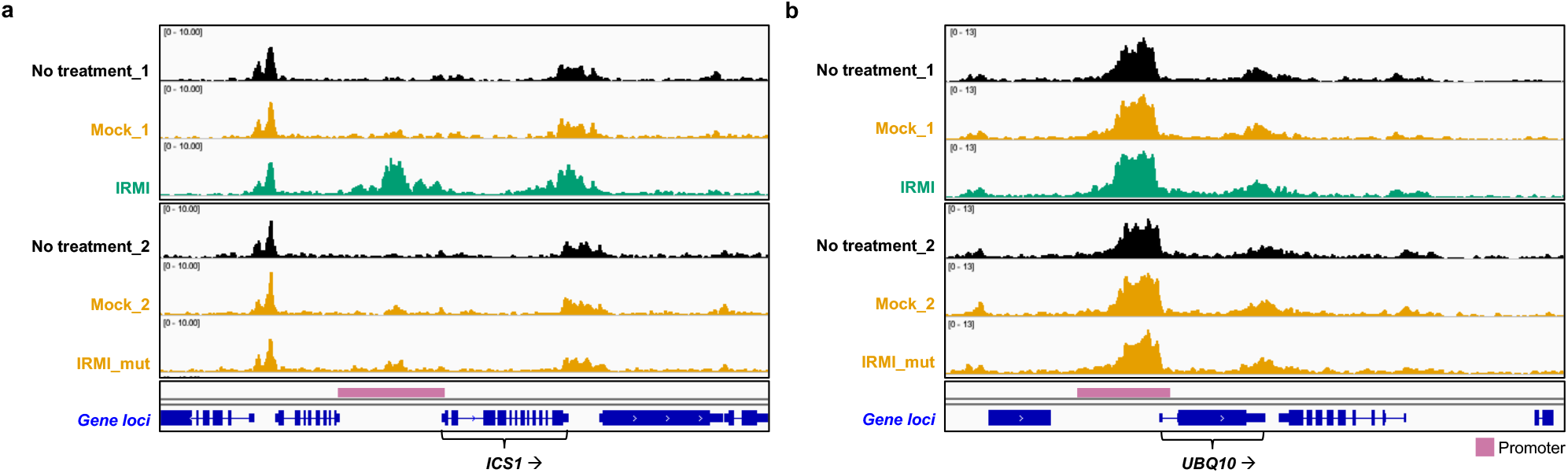
Characterization of chromatin accessible regions activated by IRMI. **(a)** and **(b)** Genome browser views of chromatin accessible regions on selected gene loci under different conditions, including IRMI. ‘No treatment_1’ and ‘No treatment_2’ are colored in black. Mock_1, Mock_2 and IRMI_mut are colored in orange. IRMI is colored in bluish green. Same color codes apply to the same corresponding conditions in the rest of this study. Gene symbols are labeled for the corresponding gene loci, (a) *ICS1* and (b) *UBQ10*. The protein coding strand is indicated in black arrow placed next to the gene symbol. The reddish-purple bars next to each gene loci indicate their putative promoter regions. **Related supplementary information:** Supplementary Fig. 8 Supplementary Tables 8

### Integration of ATAC-seq and mRNA-seq results in SRMI, IRMI and ‘SRMI+IRMI’

To identify genome-wide DARs that are activated by SRMI, IRMI and ‘SRMI+IRMI’ we normalized the ATAC peaks enriched in SRMI, IRMI and ‘SRMI+IRMI’ treatments compared to corresponding control conditions (Supplementary Fig. 9a-c). We found that DARs are upregulated at promoters of *ICS1* and *FMO1* as well as the NADPH oxidase encoding *RbohD* in response to the activation of SRMI, IRMI and ‘SRMI+IRMI’ (Fig. 4a), consistent with their upregulated gene expression in these conditions (Supplementary Fig. 6 and 7)^4^. We also found that no DARs are observed at *BIK1* locus under SRMI, IRMI and ‘SRMI+IRMI’ (Fig. 4b), though *BIK1* gene expression is induced in all these conditions (Supplementary Fig. 6 and 7)^4^. In addition, we observed that DARs at *PEP1 RECEPTOR 2* (*PEPR2*) and *SARD1* loci are only significantly induced by IRMI (Fig. 4b), but their gene expression is induced at all immune activation conditions (Supplementary Fig. 6 and 7)^4^. This indicates that not all increased DARs activated by different immune systems are positively associated with their upregulated gene expression.

**Figure 4.**
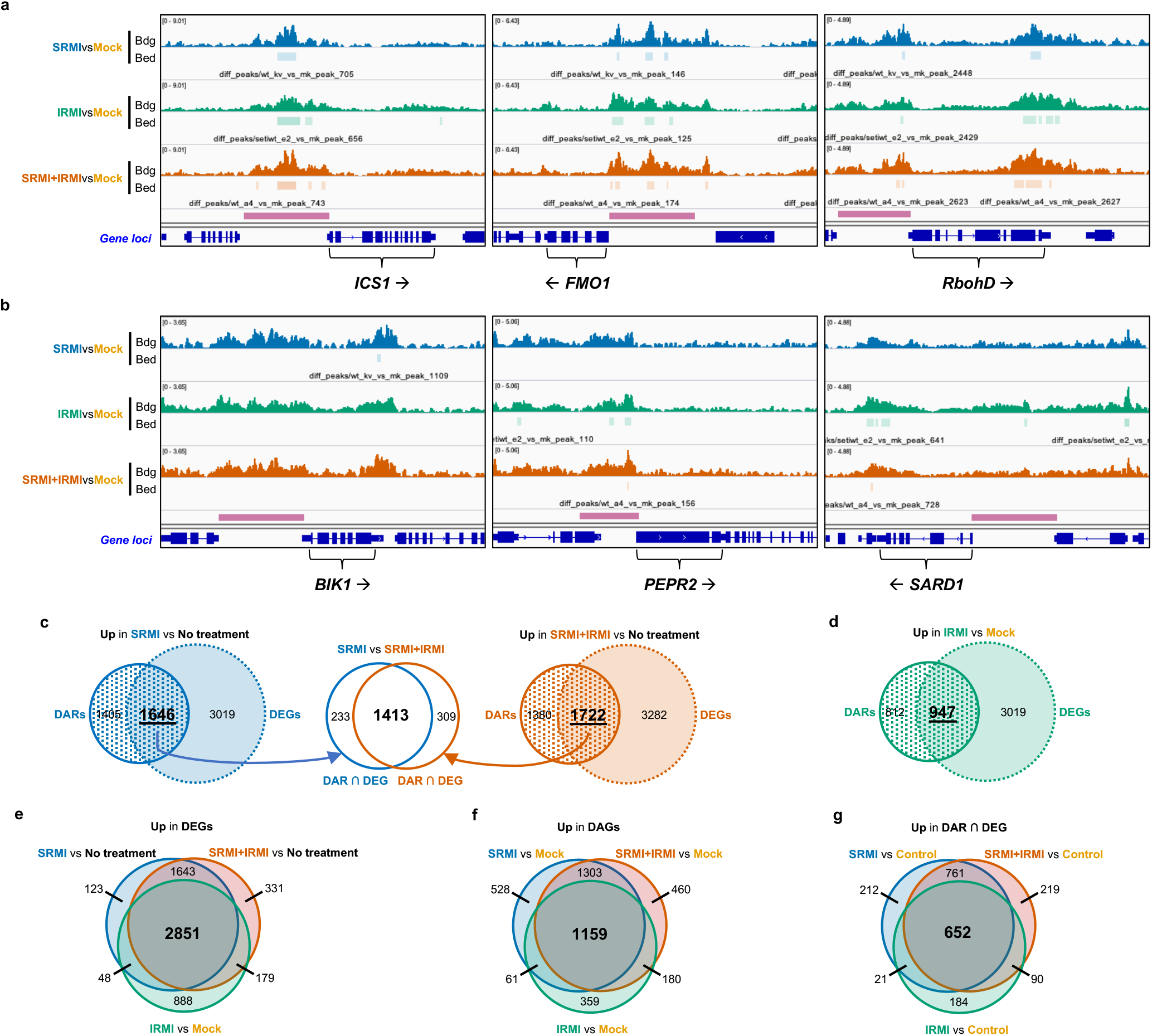
Determining chromatin accessibility changes and changes in gene expressions activated by SRMI, IRMI and ‘SRMI+IRMI’. **(a)** and **(b)** Genome browser views of differential accessible chromatin regions (ACRs) near genes that are transcriptionally upregulated in SRMI, IRMI and ‘SRMI+IRMI’ compared to Mock treatments. **(a)** Gene loci (*ICS1, FMO1* and *RbohD*) with increased ACRs in all indicated conditions. **(b)** Gene loci either have no increased ACRs in all conditions (*BIK1*) or only show increased ACRs in IRMI (*PEPR2* and *SARD1*). Other labels follow same keys indicated in Figure 1 to 3. **(c)** Integration of differential accessible regions (DAGs) and differential expression genes (DEGs) between SRMI and ‘SRMI+IRMI’. There the DAGs indicate the nearest gene loci. he Venn diagrams on the left or right sides show upregulated DARs (circle with dotted pattern) and DEGs (dashed circle with color filled) in SRMI (blue, left) or ‘SRMI+IRMI’ (vermillion, right) compared to ‘No treatment’ controls. Shared gene loci with both DARs and DEGs (intersection area in the Venn diagrams, or ‘DAR ∩ DEG’) from SRMI (n = 1635) and ‘SRMI+IRMI’ (n = 1722) were compared again in the Venn diagram in the middle. **(d)** Integration of DAGs and DEGs in IRMI. The Venn diagram (bluish green) shows upregulated DARs (circle with dotted pattern) and DEGs (dashed circle with color filled) in IRMI compared to Mock controls. **(e)** Comparisons of upregulated DEGs in SRMI, IRMI and ‘SRMI+IRMI’ compared to corresponding negative controls (‘No treatment’ or Mock). **(f)** Comparisons of upregulated DAGs in SRMI, IRMI and ‘SRMI+IRMI’ compared to corresponding negative controls (‘No treatment’ or Mock). **(g)** Comparisons of upregulated ‘DAR ∩ DEG’ in SRMI, IRMI and ‘SRMI+IRMI’ compared to Controls (‘No treatment’ or Mock). **Related supplementary information:** Supplementary Fig. 7, 8 and 9 Supplementary Tables 7, 8 and 9

To determine the extent to which activated open chromatin regions from the ATAC-seq analysis are correlated with induced gene expression in all immune conditions we integrated our ATAC-seq data with corresponding mRNA-seq data. We found 1646 gene loci with increased ATAC peaks (DARs) as well as significantly upregulated gene expression (DEGs) in SRMI versus ‘No treatment’, and 1722 such loci in ‘SRMI+IRMI’ versus ‘No treatment’ (Fig. 4c; Supplementary Tables 9). By comparing the intersection of the positively correlated gene loci (‘DAR ∩ DEG’), we found substantial overlap (1413 gene loci) between these two conditions (Fig. 4c; Supplementary Tables 9). Comparing ‘IRMI vs Mock’, we found 947 loci showing positive correlation of increased DARs and upregulated DEGs (Fig. 4d; Supplementary Tables 9). The same GO terms are enriched in both 1413 and 947 loci lists (Supplementary Fig. 9c,d; Supplementary Tables 9). Thus, a common set of genes is activated during SRMI, IRMI and ‘SRMI+IRMI’, and transcriptional activation might require chromatin in these gene loci to open up for active transcription events.

To better understand DEGs and DAGs that are induced by SRMI, IRMI and ‘SRMI+IRMI’, we individually compared upregulated DEGs, DAGs and ‘DAR∩DEG’ that are activated in these conditions compared to corresponding control conditions (Fig.4 e-g; Supplementary Tables 9). We found a large proportion of both upregulated genes and increased DAGs are shared by all three immune activation conditions (Fig. 4e,f; Supplementary Tables 9). We then compared SRMI, IRMI and ‘SRMI+IRMI’ upregulated DEGs and DARs (‘DAR∩DEG’), and identified 782 gene loci are shared by SRMI, IRMI and ‘SRMI+IRMI’ (Fig. 4g, Supplementary Tables 9). These responses shared by SRMI, IRMI and ‘SRMI+IRMI’ could reveal common transcriptional regulatory mechanism, where a common set of TFs might be required for controlling gene expression.

### Transcriptional gene regulatory networks of SRMI, IRMI and ‘SRMI+IRMI’

Identification of ACRs can assist to determine locations of putative CREs, where transcriptional regulators, especially DNA-binding proteins such as TFs, might bind. To identify regulatory interactions between TF regulators and target genes, gene regulatory networks (GRNs) were delineated through the integration of RNA-Seq, ATAC-Seq and TF motif information^47^. GRNs at an early time point (4 hpi) upon activation of SRMI, IRMI and ‘SRMI+IRMI’ were constructed, based on motifs enriched for DAR in these conditions. TF binding site mapping data for 1,793 motifs, corresponding to 916 Arabidopsis TFs were used to link specific regulators with putative target genes, based on the motif location in the DAR and the closest gene^47^. To narrow down the list of TFs, we selected those which showed increased gene expression (log2FC > 1, q-value < 0.01, Supplementary Tables 10). We identified 115, 34 and 133 TFs as regulators in SRMI, IRMI and ‘SRMI+IRMI’, based on the significant enrichment of 210, 73 and 248 motifs in the corresponding DARs (Fig. 5a, Supplementary Tables 10). Comparing regulators between the different conditions reveals that 25 regulators, of which 72% are WRKY TFs, are common to all 3 networks, while 82 regulators are shared between SRMI and ‘SRMI+IRMI’, corresponding predominantly with WRKY, bHLH and bZIP TFs (Fig. 5b). This result reveals a diversity of TF families is playing an important role in the transcriptional reprogramming of gene expression during the activation of plant immunity.

To assess the biological processes controlled by these different regulators, GO enrichment was performed on each set of target genes, per network and per TF. We found ‘response to chitin’, ‘response to bacterium’ and ‘response to hypoxia’ are the top three GO terms that are commonly enriched in SRMI, IRMI and ‘SRMI+IRMI’ (Fig. 5c, Supplementary Fig. 10, Supplementary Tables 10). Whereas most WRKYs are activated during immune responses independently of whether the activation occurs through surface or intracellular receptors, WRKY65 and WRKY59 are specific to IRMI, and the targets of WRKY59 are enriched in ‘regulation of cell death’ (Fig. 6, Supplementary Fig. 10b). Some examples of TFs implicated by this analysis, some of which have been confirmed in controlling hormone-related processes, are: (1) response to JA and SA in SRMI: AtIDD5; (2) response to SA in SRMI: KAN2, WRKY33, WRKY45, TGA7, and JKD; (3) SA signaling in IRMI: AT5G01380; and (4) response to SA in ‘SRMI+IRMI’: KAN2, ANAC029, ANAC046, ABO3, TGA3 and TGA7 (Supplementary Fig. 10). Response to ABA was only observed for 8 WRKY TFs in SRMI (WRKY7, 11, 15, 17, 22, 40, 45 and 75), which might be mostly associated with the wounding response. Stronger induction of IRMI in addition to SRMI might be more dominant to this ABA or wounding-associated transcriptional regulation. Two members of the CAMTA transcription factor family (CAMTA1 and CAMTA3) are exclusively enriched in IRMI and were previously characterized as repressors of SA-regulated genes. However, upon pathogen infection CAMTA-mediated repression is alleviated and plant defense genes are expressed^48–50^. These results indicate that the function of these CAMTA transcription factors involved in immunity are mediated by intracellular receptors. For SRMI there is only one TF exclusive to this condition, CBF2, that regulates a SRMI-specific GO term, ‘toxin metabolic process’.

**Figure 5.**
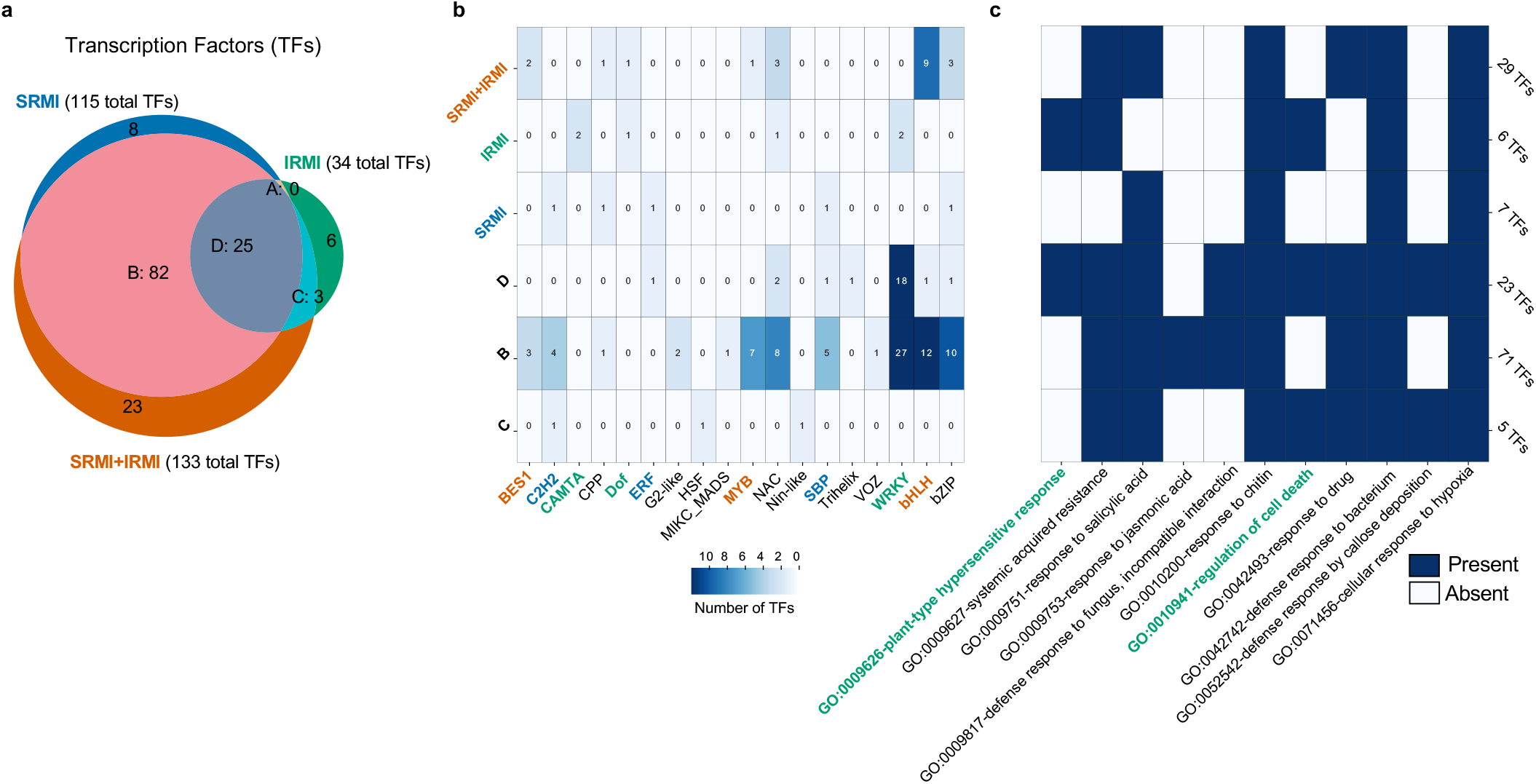
Overview of regulatory TFs across the different gene regulatory networks. **(a)** Venn diagram showing shared and unique TFs across the different networks inferred for each condition. **(b)** Heatmap with the number of TF family members in each of the Venn diagram sets, as indicated by the coloring of each cell. TFs that are specific to SRMI, IRMI and ‘SRMI+IRMI’ are highlighted with corresponding color code. **(c)** Heatmap displaying presence or absence of GO terms enrichment in each of the Venn diagram sets. Numbers at the right indicate the number of TFs showing GO enrichment. TFs that are specific to IRMI are enriched in ‘plant-type hypersensitive response’ and ‘regulation of cell death’ (in blueish green). **Related supplementary information:** Supplementary Fig. 10 and 11 Supplementary Tables 10

**Figure 6.**
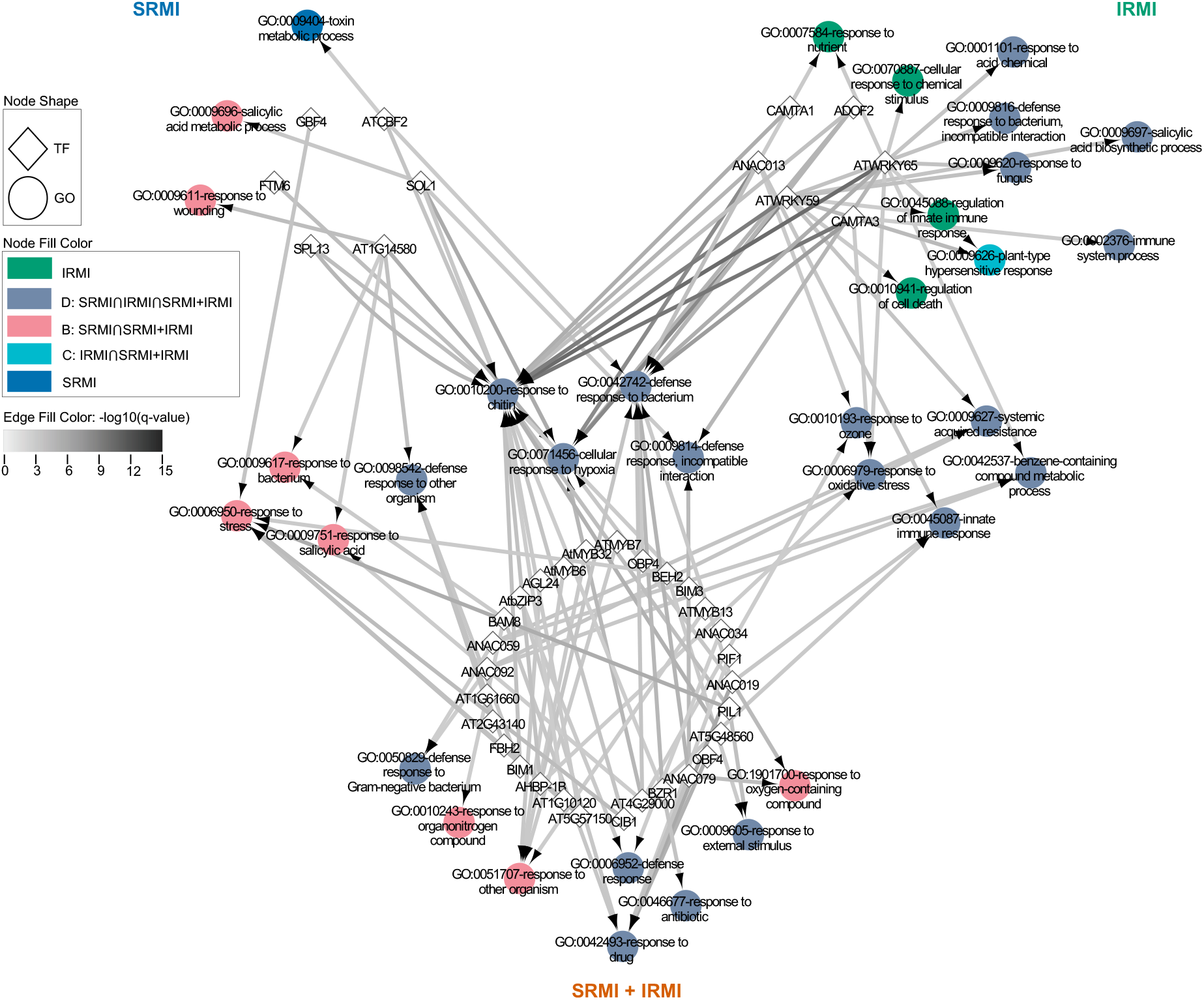
TF-GO term network for condition-specific TFs. Network diagram with the TFs specific for each condition ‘SMRI’, ‘IRMI’ and ‘SRMI+IRMI’ organized in circles linked to the GO terms enriched in the putative target genes they control. The GO terms are colored according to whether they are exclusive of any specific condition or if they belong in any of the possible intersections of the conditions. The GO terms controlled by TFs exclusive of one condition are in the most outer parts. The GO terms controlled by TFs exclusive of two and three conditions are located closer to the center and in the center, respectively. Note there are not GO terms shared between TFs of SRMI and IRMI. Also, there are no GO terms that are exclusive of ‘SRMI+IRMI’ and ‘SRMI ⋂ IRMI’ because none of them are controlled by SRMI, IRMI or ‘SRMI+IRMI’ exclusive TFs. **Related supplementary information:** Supplementary Fig. 12 Supplementary Tables 10

The clustering coefficient of a network is a measure of the tendency of the nodes to cluster together, which for GRNs indicates that for specific genes, the incoming regulatory TFs are also controlling each other, suggesting TF crosstalk. The clustering coefficient is significantly higher for SRMI and IRMI networks than for ‘SRMI+IRMI’ and the intersection of SRMI, IRMI and ‘SRMI+IRMI’ (SRMI∩IRMI∩’SRMI+IRMI’, or ‘Set D’) (Supplementary Fig. 11). The IRMI network stands out having 23 genes with a clustering coefficient of 0.5 or higher, which are controlled by a combination of WRKY TFs (WRKY6, 26, 31, 40, 47 and 70), CAMTA3, HSFB2A and IDD1 (Supplementary Fig. 12). Overall, the inferred networks revealed that both shared and unique regulators are involved in controlling gene expression, with an important role for WRKY TFs controlling the ‘response to bacteria’ as well as other TFs regulating other hormone-related biological processes. In addition, the tight control of specific target genes by multiple TFs, some also controlling each other, enables investigation into the hierarchy of TF signaling in different types of immunity activation.

## Discussion

Our understanding of transcriptional regulation in eukaryotes has been greatly advanced by sequencing methods developed for chromatin biology^51^. Genome-wide chromatin accessibility data for different plant species have demonstrated interesting aspects of mechanisms involved in transcriptional regulation of diverse biological processes^20–23,37,52^, but rarely for plant immunity. Several TFs have been implicated in plant innate immunity^11,53^. How these TFs function in a regulatory network has remained poorly understood. Here we report chromatin accessibility landscapes that are activated by both cell-surface- and intracellular-immune receptor-mediated immunity (SRMI and IRMI). There are few studies of IRMI in the absence of SRMI, and we highlight here the similarity and differences for changes in chromatin accessibility and associated gene expression between these two systems.

From the minimum GRNs we constructed here based on our ATAC-seq and RNA-seq results, we show that WRKY transcription factors are the predominant players in the GRNs regulating most genes that are activated during both SRMI and IRMI. However, due to incomplete public data, our GRNs here cannot reflect all regulatory possibilities. For instance, the DNA-binding motifs of some TFs are still not known^54^. In addition, we prioritized upregulated genes in our analysis, and negative regulation of some genes upon TF binding might also play an important role. These networks will be further improved when more data become available.

For upregulated DEGs that show increased DARs, the binding of TFs might be correlated with the chromatin opening. However, it is challenging to distinguish whether TF binding is cause or consequence of chromatin opening. Some TFs can serve as ‘pioneer’ TFs to initiate transcription by recruiting ‘non-pioneer’ TFs, other transcriptional regulators and the RNA polymerase II machinery^51,55,56^. Such ‘pioneer’ TFs may recruit components that can open the chromatin, such as histone remodelers^51^. However more genetic evidence is required to evaluate this hypothesis.

The chromatin accessibility landscapes and implied GRNs reported here provide a snapshot of events during the activation of different immune responses. Transcription, like many other processes, is a dynamic, so it is important to profile the changes in chromatin accessibilities and corresponding gene expressions over a time-course. With time-series ATAC-seq and RNA-seq data in future studies, we will be able to generate dynamic transcriptional regulatory networks that will provide more novel insights into transcriptional regulatory mechanisms required for immune activation and for the establishment of disease resistance.

## Methods

### Plant materials and growth condition

*Arabidopsis thaliana* Columbia-0 (Col-0) accession was used as wild type in this study. SETI^WT^ and SETI^mut^ transgenic plants used have been described previously^9^. Seeds were sown on compost and plants were grown at 21 °C with 10 hours under light and 14 hours in dark, and at 70% humidity. The light level is approximately 180-200 µmols with fluorescent tubes.

### Activation of SRMI and ‘SRMI+IRMI’

*Pseudomonas fluorescens* engineered with a type III secretion system (Pf0-1 ‘EtHAn’ strains) expressing one of wild-type AvrRps4 or AvrRps4 KRVY135-138AAAA mutant effectors were grown on selective KB plates for 24 h at 28 °C^6,35^. Bacteria were harvested from the plates, resuspended in infiltration buffer (10 mM MgCl_2_) and the concentration was adjusted to OD_600_ = 0.2 (10^8^ CFU•mL^-1^). The abaxial surfaces of 5-week-old Arabidopsis leaves were hand infiltrated with a 1-mL disposable needleless syringe (Slaughter Ltd, R & L, catalogue number: BS01T). Leaves infiltrated with 10 mM MgCl_2_ serves as mock treatment. Leaves infiltrated with Pf0-1:AvrRps4^WT^ activates ‘SRMI+IRMI’, and those infiltrated with Pf0-1:AvrRps4^mut^ activates SRMI only^4^.

### Activation of IRMI

5-week-old Arabidopsis leaves of SETI^WT^ (E2:AvrRps4^WT^) infiltrated with 50 µM β-estradiol (E2, Sigma-Aldrich, catalogue number: E8875; dissolved in 100% dimethyl sulfoxide, also known as DMSO, Sigma-Aldrich, catalogue number: D8418) activates ‘IRMI’ only, as described previously^9^. 0.1% DMSO (same titrate as 50 mM E2 stock solution diluted in pure water and generating 50 µM E2 working solution) in pure water used as mock treatment for infiltration with a 1-mL needleless syringe. SETI^mut^ (E2:AvrRps4^mut^) with similar treatments as in SETI^WT^ serve as additional negative IRMI controls, as described previously^9^.

### RNA isolation and sequencing (RNA-seq)

Leaf samples from SRMI, ‘SRMI+IRMI’, IRMI were isolated as described previously^8^. Total RNA samples are sent by dry ice to BGI for mRNA isolation and library construction and sequenced on BGISEQ-500 sequencing platforms.

### RNA-seq raw data processing, alignment and quantification of expression and data visualization

Raw reads are trimmed into clean reads by BGI bioinformatic service into 50 bp. At least 12 million single-end clean reads for each sample were provided by BGI for RNA-seq analysis. All reads have passed FastQC before the following analyses (Simon Andrews; FastQC: https://www.bioinformatics.babraham.ac.uk/projects/fastqc/). All clean reads are either mapped to TAIR10 Arabidopsis genome/transcriptome via TopHat2 or to a comprehensive Reference Transcript Dataset for Arabidopsis (AtRTD2) containing 82,190 non-redundant transcripts from 34,212 genes via Kallisto (SRMI and ‘SRMI+IRMI’) or Salmon (IRMI) tools^57–60^. Detailed scripts and versions of each software can be found via the GitHub link: https://github.com/TeamMacLean/fastqc_kallisto_analysis. Mapped reads were sorted with SAMtools and visualized in Integrative Genomics Viewer (IGV) with TAIR10 reference genome^57,61,62^. The estimated gene transcript counts were used for differential gene expression analysis, statistical analysis and data through the 3D RNA-seq software^63^. For both datasets, the low expressed transcripts were filtered if they did not meet the criteria ≥ 3 samples with ≥ 1 count per million reads (CPMs). An expressed gene must have at least one expressed transcript. The batch effects between biological replicates were removed to reduce artificial variance with RUVSeq method^64^. The expression data was normalized across samples with TMM method^65^. The significance of expression changes in the contrast groups ‘SRMI vs no treatment’ and ‘SRMI+IRMI vs no treatment’, and ‘IRMI vs Control_1’ and ‘IRMI vs IRMI_mut’ were determined by the limma-voom method^66,67^. A gene was defined as significant DEG if the BH adjusted p-value < 0.01 and log_2_(fold_change) ≥ 1.

### FANS-ATAC-seq

Leaf samples from SRMI, ‘SRMI+IRMI’, IRMI and control conditions were collected at 4 hpi of treatment (same time points as RNA-seq samples) or without any treatment. 2 fully expanded leaves with treatment from each plant and 3 plants in total are collected as one sample and one biological replicate. 6 leaves of one sample were chopped quickly in 1 mL 4-°C-prechilled phosphate-buffered saline (PBS) buffer (1×, pH 7.4) with a clean unused razor blade (Agar Scientific Ltd, catalogue number: T586) into fine pieces (less than 1 min). The leaf lysis containing crude nuclei extract was transferred and filtered through a 30-µm CellTrics^®^ cell strainer (Sysmex, catalogue number: 04-0042-2316) into a 100×16-mm round-base test tube (Slaughter Ltd, R & L, catalogue number: 142AS) with 1-mL sterile tip by pipetting. The sharp end of the tip was cut off and shortened with 2-mm in length by a pair of sterile scissors to minimize the damage to the nuclei. All samples of leaf lysis were kept on ice immediately after the transfer. 1-mL CyStain PI Absolute P nuclei 4’,6-diamidino-2-phenylindole (DAPI) staining buffer (Sysmex, catalogue number: 05-5022) was added into each nuclei extract. Nuclei extract with the staining buffer were gently mix and kept on ice. Stained nuclei extract were submitted to BD FACSMelody Cell Sorter with a green laser for fluorescence-assisted nuclei sorting (FANS) with a similar setting as described previously^19^. FANS-purified nuclei samples were collected in 1.5 DNA LoBind Eppendorf microcentrifuge tubes (Fisher Scientific, catalogue number: 10051232) and kept on ice. Nuclei pellets were collected as described previously by centrifugation at 1,000 × *g* and tagmented with Nextera DNA Library Prep Kit (Illumina, catalog number: FC-121-1030, now discontinued; replacement can be found as Illumina Tagment DNA TDE1 Enzyme and Buffer Kits, catalog numbers: 20034197 or 20034198)^68^. We used 0.1 µL TDE1 enzyme in a 5 µl total reaction mix for each ATAC samples. The following PCR library construction and quality control steps were performed as recommended^68^. The only difference here was that we used dual index primers designed by ourselves for barcoding the libraries and multiplexing. Those primers have been validated in our previous experiments^8^, and the detailed sequence information can be found in Supplementary Table 1. Multiplexed ATAC-seq libraries were sequenced with multiple NextSeq 500/550 High Output Kits (75 Cycles) on an in-house NextSeq 550 sequencer.

### ATAC-seq raw data processing and alignment

Sequencing results were demultiplexed using bcl2fastq tool to generate adaptor-trimmed raw reads. Pair-end and 37 bp each end reads were tested with FastQC and Picard tools for quality control and testing PCR duplications. Raw reads were mapped to TAIR10 Arabidopsis reference genome with Bowtie2 and sorted with SAMtools^61,69^. Reads mapped to chloroplast and mitochondria were removed before the follow-up analyses. Detailed scripts, software versions and QC outputs can be found in the GitHub link: https://github.com/TeamMacLean/dingp_atacseq_for_publication.

### Identification of ACRs

Identification of ACRs was done by callpeak function in MACS v.2.2.7^70^. All replicates of samples under specific condition were used as input of treatment and genomic DNA samples were used as input of control. For visualization of fold enrichment of mapped reads compared control samples, FE.bdg files were generated by bdgcmp function in MACS. FE.bdg files were visualized by IGV^62^. In the trial analysis of FANS-ATAC-seq, we counted mapped reads for ACRs using coverage function in Bedtools v.2.28.0^71^. Then, we made correlation plots based on log2 read counts of each common ACR between replicates to find out the reproducibility for 10k, 20k, 50k, and 80k samples using our R script, which is listed in our GitHub link: https://github.com/slt666666/Plant_Innate_Immunity_ATAC-seq.

### Profile of ACRs binding to TSS/TTS regions

The heatmap of ACRs binding to TSS regions and the distribution of ACRs binding to TSS and TTS regions were obtained by ChIPSeeker v.1.24.0 package within R^72^. The features of ACRs were annotated by ChIPSeeker package using the annotatePeak function. In this part, promoters were defined as −2,000 to 1,000 bp from the TSS. The Upset plot which showed ACRs shared on several conditions were generated by UpSetR package based on the nearest genes from ACRs^73^.

### Identification of DARs

Identification of DARs is achieved by applying callpeak function of MACS. All replicates of samples under specific condition were used as input of treatment and all replicates of samples under corresponding control conditions were used as input of control. Annotation of genes with enriched DARs within 2 kb upstream of a gene is done by annotatePeakInBatch function in ChIPpeakAnno package^72^. Annotation of genes with the other DARs in distal intergenic genome loci is done by our Python script.

### Integration of DEGs and DARs

The identification of common genes to annotated genes with enriched DARs and significantly upregulated (log2FC > 1, q-value < 0.01) genes is done by our Python script. The GO analysis for these common genes is conducted by g:Profiler^74^. All scripts used for the analyses of ACRs and integration of DEGs and DARs are available: https://github.com/slt666666/Plant_Innate_Immunity_ATAC-seq.

### Motif-based inference of gene regulatory networks using ACRs

The inference of GRNs was done using an ensemble motif mapping method described previously^75^, combining all the matches from Find Individual Motif Occurrences (FIMO) with the top 7000 matches from Cluster-Buster (CB)^76,77^. This mapping information was used to determine which motifs were significantly enriched in the ACRs derived from the ATAC-seq experiments for each condition (SRMI, IRMI and ‘SRMI+IRMI’). Per condition, for TFs showing differential expression the associated motifs were tested for enrichment in the ACRs and significant motifs were retained (adjusted p-value ≤ 0.01). Based on motif coordinates in ACRs, individual motif matches were assigned to the closest gene, establishing a link between the TFs that bind these motifs and putative target genes. The differential expression information was integrated to identify only the TFs and target genes that were differentially expressed for each condition. Finally, for each TF, the putative target genes set was analyzed for over-represented GO Biological Process terms (only using experimentally and curated annotations; hypergeometric distribution q-value < 0.001).

## Data availability

All raw reads in this study have been uploaded to the European Nucleotide Archive (ENA) as data repository and can be retrieved through accession number PRJEB34955 and PRJEB38924 for RNA-seq and PRJEB38923 for ATAC-seq. For data reproducibility, all scripts generated in this study, software versions can be found in related GitHub links indicated in the Methods section.

## Acknowledgements

We thank the Gatsby Foundation (United Kingdom) for funding to the JDGJ laboratory. PD acknowledges support from the European Union’s Horizon 2020 Research and Innovation Program under Marie Skłodowska-Curie Actions (grant agreement: 656243) and a Future Leader Fellowship from the Biotechnology and Biological Sciences Research Council (BBSRC) (grant agreement: BB/R012172/1). TS, RKS, DM and JDGJ were supported by the Gatsby Foundation funding to the Sainsbury Laboratory. NMP and KV were supported by a BOF grant from Ghent University (grant agreement: BOF24Y2019001901). WG and RZ were supported by Scottish Government Rural and Environment Science and Analytical Services division (RESAS), and RZ also acknowledge the support from a BBSRC Bioinformatics and Biological Resources Fund (grant agreement: BB/S020160/1). BN was supported by the Norwich Research Park (NRP) Biosciences Doctoral Training Partnership (DTP) funded by BBSRC (grant agreement: BB/M011216/1). SH and XF were supported by a BBSRC Responsive Mode grant (grant agreement: BB/S009620/1) and a European Research Council Starting grant ‘SexMeth’ (grant agreement: 804981). CL was supported by Deutsche Forschungsgemeinschaft (grant agreement: LI 2862/4).

## Author contributions

PD and JDGJ conceptualized and designed the research. PD conducted all experiments with assistance from BPMN and SH. PD together with TS, RKS, NMP, WG and CL performed data analyses with assistance from DM, RZ and KV. PD wrote the manuscript with input from all co-authors. All authors helped editing and finalizing the manuscript.

## Supplementary information

**Supplementary Figure 1.**
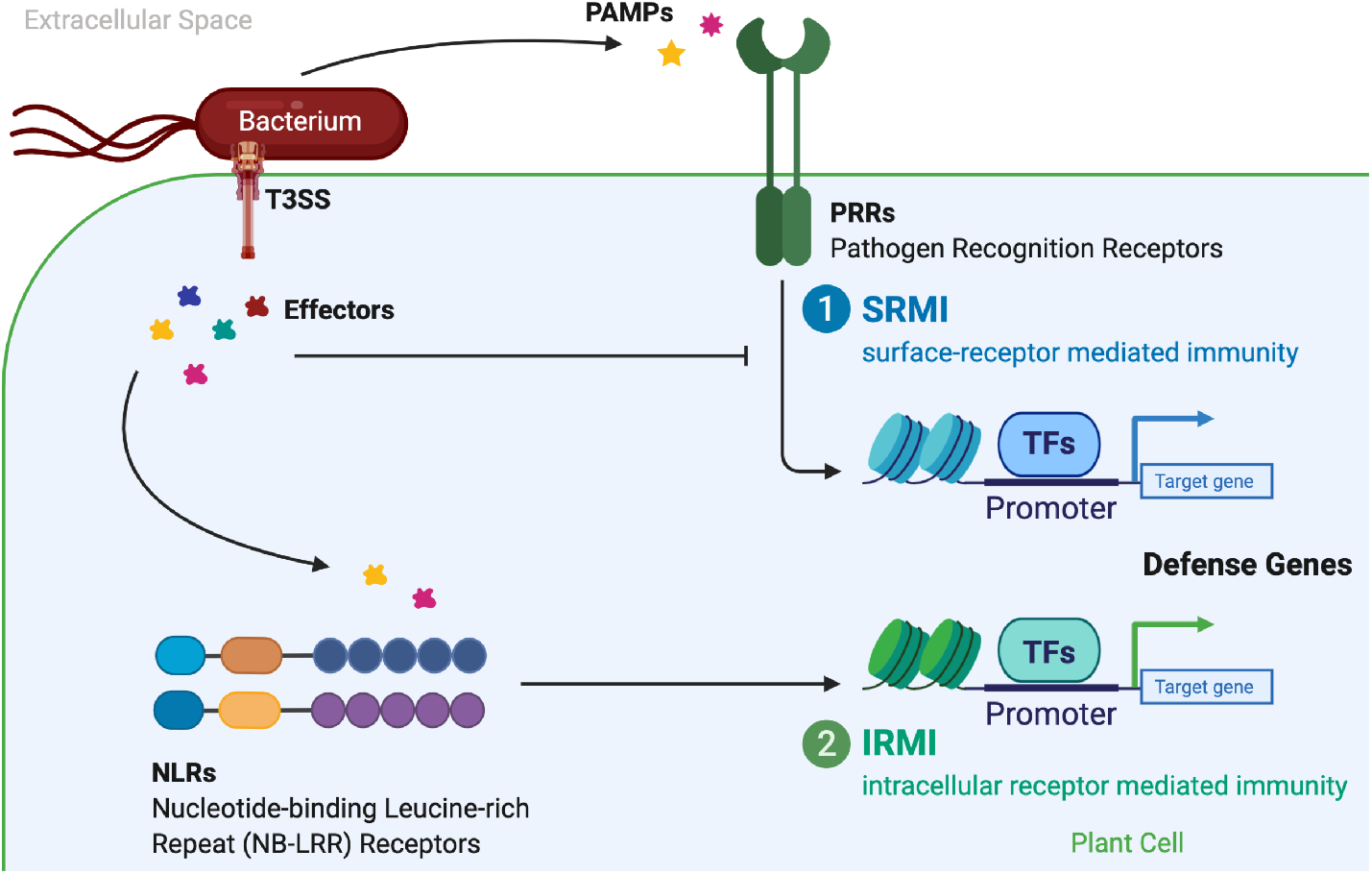
Schematic view of integrated plant innate immune system. Here plant-pathogen interactions are reflected by plant-bacteria interactions. Bacterial plant pathogens propagate exclusively in the extracellular spaces of plant issues. Pathogen-Associated Molecular Patterns (PAMPs) released from the pathogens into the extracellular spaces, such as flagellin and elongation factor thermo unstable (EF-Tu) are recognized by cell surface Pattern Recognition Receptors (PRRs) and elicit Pattern-Triggered Immunity (PTI) or here as a refined term cell Surface Receptor-Mediated Immunity (SRMI). Bacterial pathogens deliver effector proteins into the host cell by a type-III secretion pilus system (T3SS). These intracellular effectors often act to suppress SRMI. However, many are recognized by intracellular nucleotide-binding (NB)-LRR receptors (NLRs), which induces Effector-Triggered Immunity (ETI) or here as a refined term Intracellular Receptor-Mediated Immunity (IRMI). Both SRMI and IRMI can lead to the transcriptional reprogramming by recruiting transcription factors (TFs) to the target gene loci, such as their promoters, and activate defense gene expression for effective bacterial resistance.

**Supplementary Figure 2.**
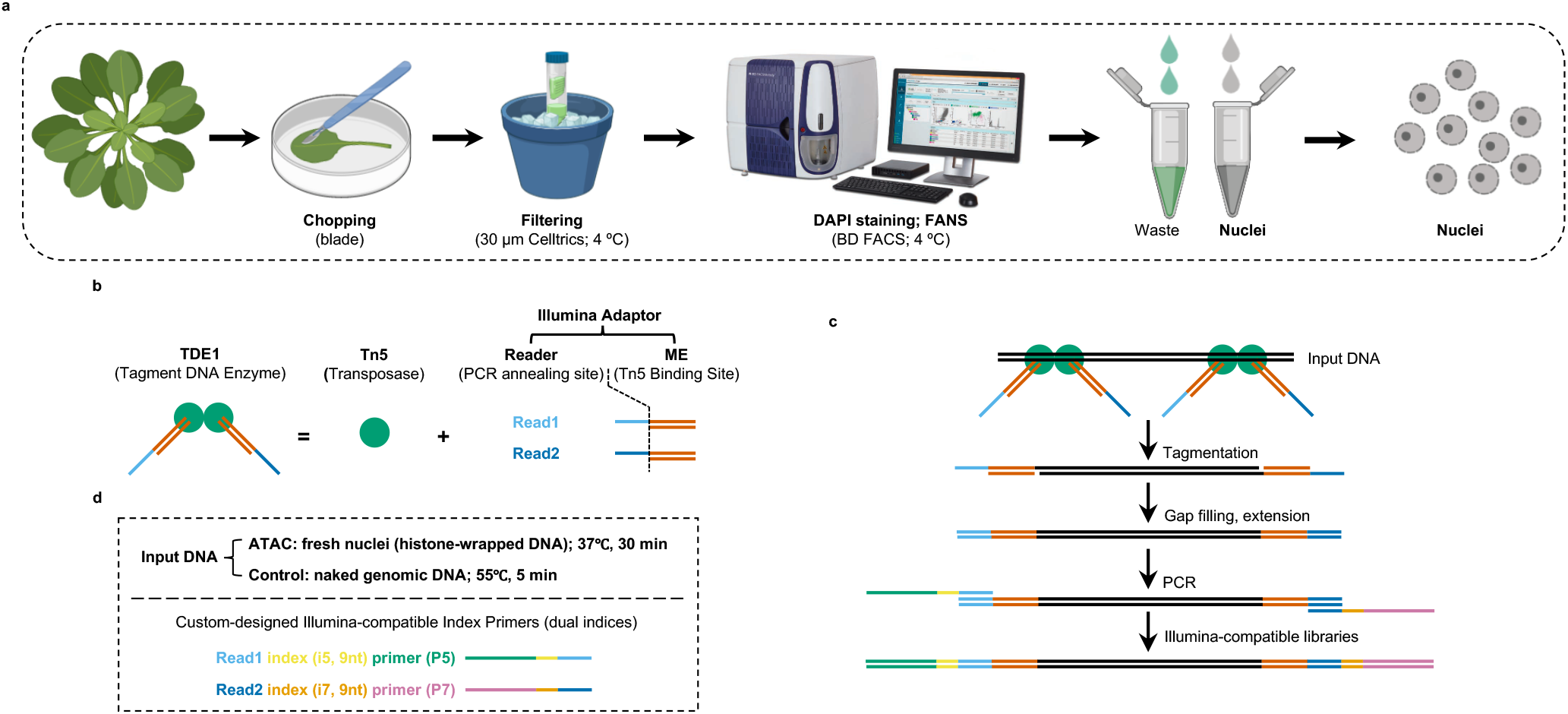
Establishment of FANS-ATAC-seq for Arabidopsis leaf tissue. **(a)** A simplified workflow for fluorescence-assisted nuclei sorting (FANS) of leaf nuclei purification. **(b)** Composition of Tn5 transposasome for DNA tagmentation. **(c)** Schematic view of assay for transposase-accessible chromatin followed by sequencing (ATAC-seq).

**Supplementary Figure 3.**
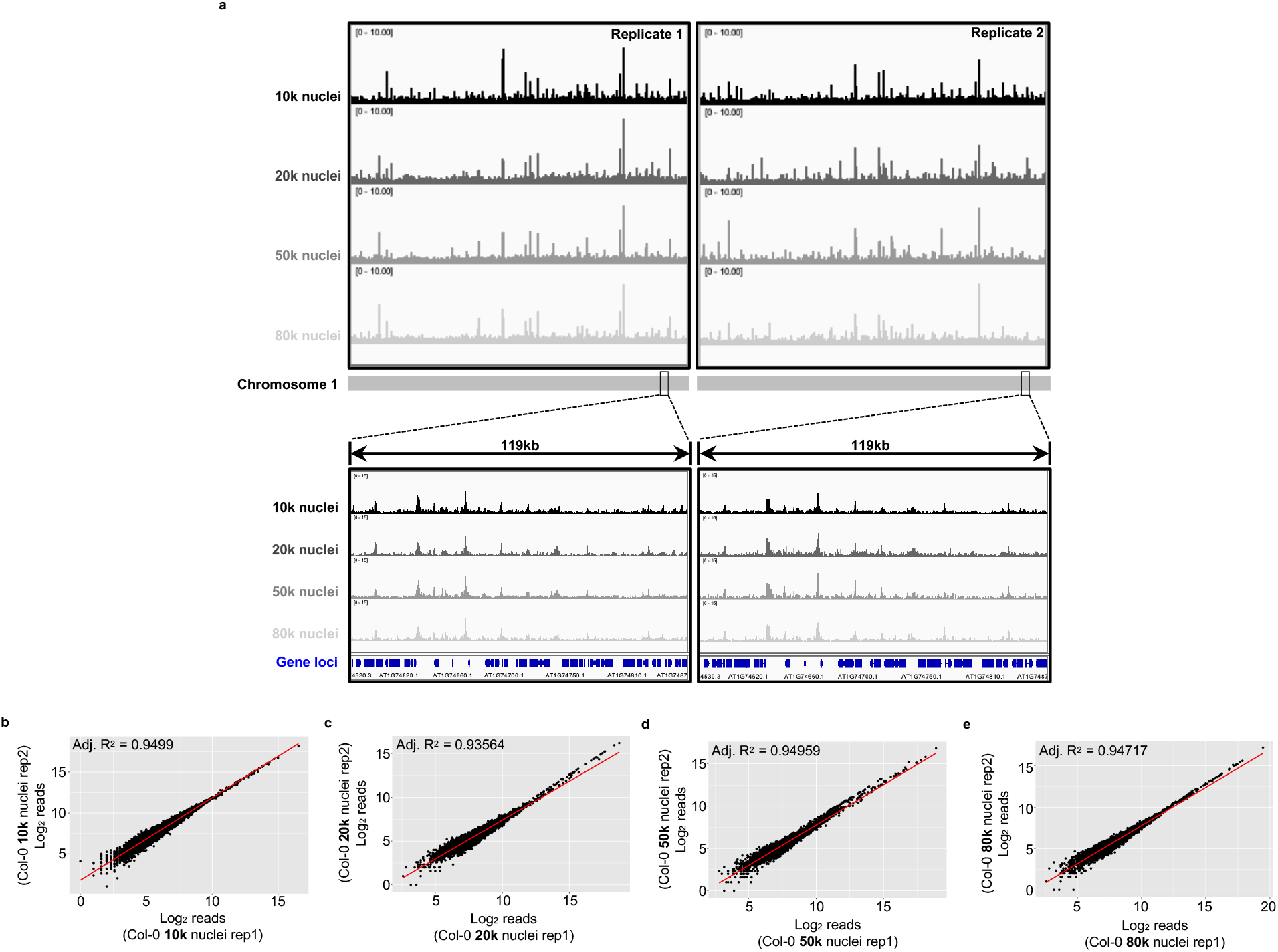
Trial run results of FANS-ATAC-seq for Arabidopsis leaf tissue demonstrate a good reproducibility. **(a)** Snapshot of FANS-ATAC-seq results on the Integrative Genomics Viewer (IGV) genome browser from the trial experiment. **(b-e)** Correlation co-efficiency analysis for different leaf nuclei input in FANS-ATAC-seq trial experiment. **(a), (b), (c)** and **(d)** are analysis between the first biological replicate (replicate 1 or rep1) and rep 2 for 10,000 (10k), 20k, 50k and 80k nuclei input, respectively. Adjusted R square (R^2^) values from all individual comparisons are all greater than 0.9, indicating a high reproducibility of the FANS-ATAC-seq results from FANS-ATAC-seq workflow established in this study.

**Supplementary Figure 4.**
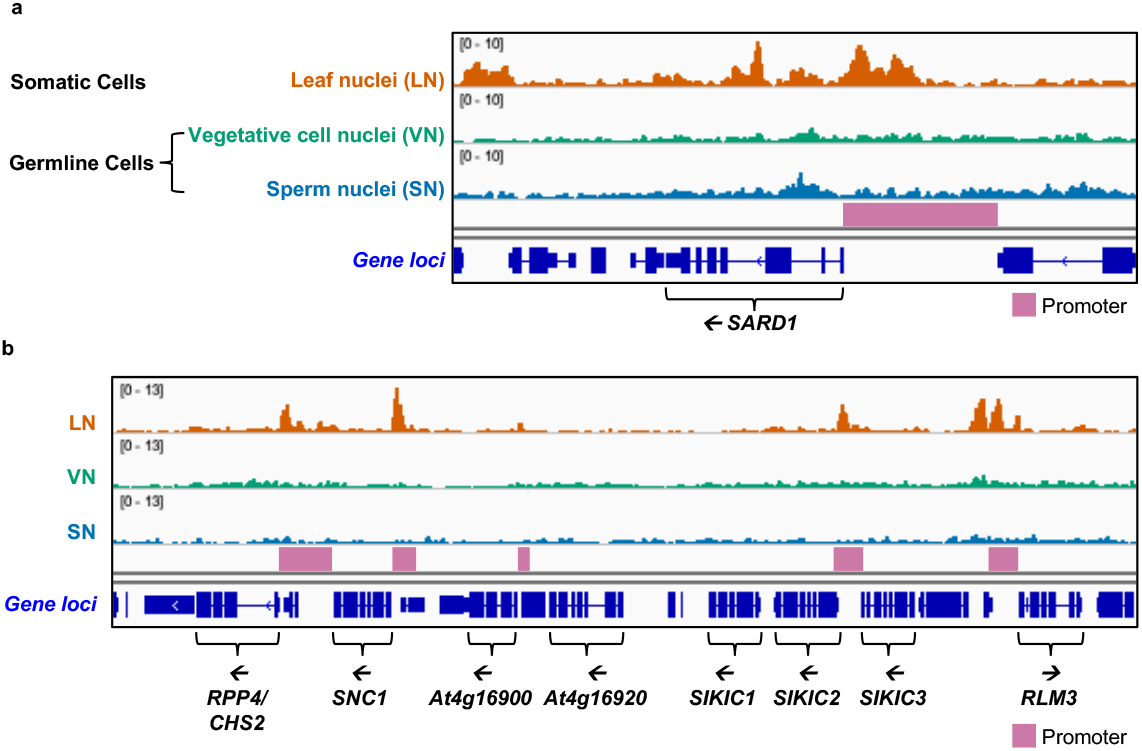
FANS-ATAC-seq using Arabidopsis leaf nuclei, vegetative and sperm nuclei derived from Arabidopsis pollen grain show tissue-specific chromatin accessibility. **(a)** Comparative view of accessible chromatin regions (ACRs) between somatic and germline cells at *SARD1* gene locus. **(b)** Comparative view of ACRs between somatic and germline cells at a *Resistance*(*R*)-gene NLR gene cluster loci.

**Supplementary Figure 5.**
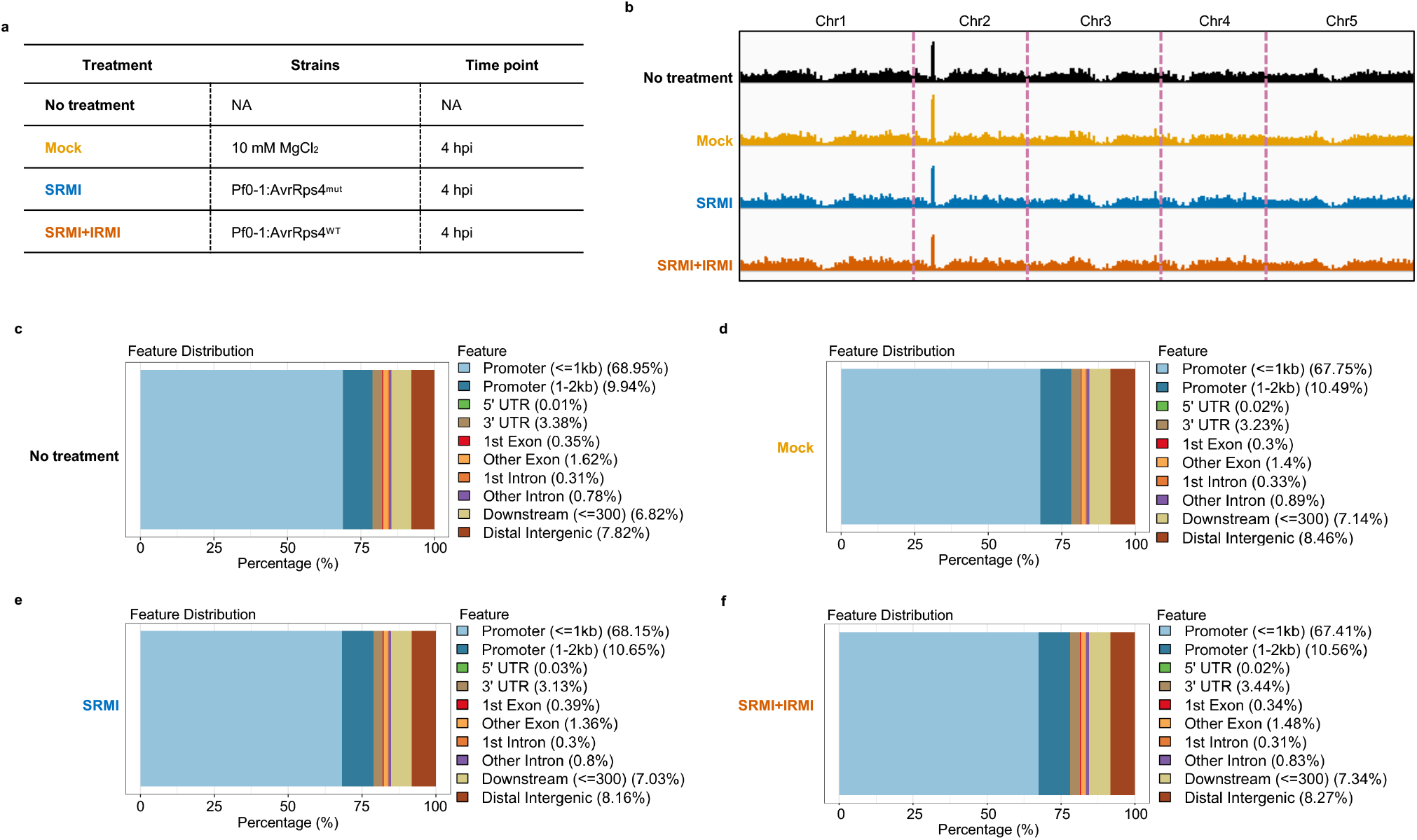
Additional information of FANS-ATAC-seq for chromatin landscapes activated by SRMI and ‘SRMI+IRMI’. **(a)** Simplified table for the information of experimental design for RNA-seq. Four conditions include SRMI, ‘SRMI+IRMI’ with ‘No treatment’ and Mock as control. SRMI is implied by the infiltration of Pf0-1 EtHAn carrying mutant AvrRps4 (KRVY 135-138 to AAAA) (Pf0-1:AvrRps4^mut^), which cannot activate NLRs. ‘SRMI+IRMI’ is activated by the infiltration of Pf0-1:AvrRps4^WT^ that is recognized by NLRs RRS1/RPS4 and RRS1B/RPS4B. Mock infiltration with 10 mM MgCl_2_ serves as control for wounding induced during the infiltration process. All sample with infiltration were collected at 4 hours post infiltration (hpi) and immediately used for FANS-ATAC-seq. **(b)** Genome browser views of ATAC-seq-indicated chromatin accessibility genome-wide coverage under different conditions. **(c-f)** Accessible chromatin regions from all conditions are mapped to different genomic features as indicated.

**Supplementary Figure 6.**
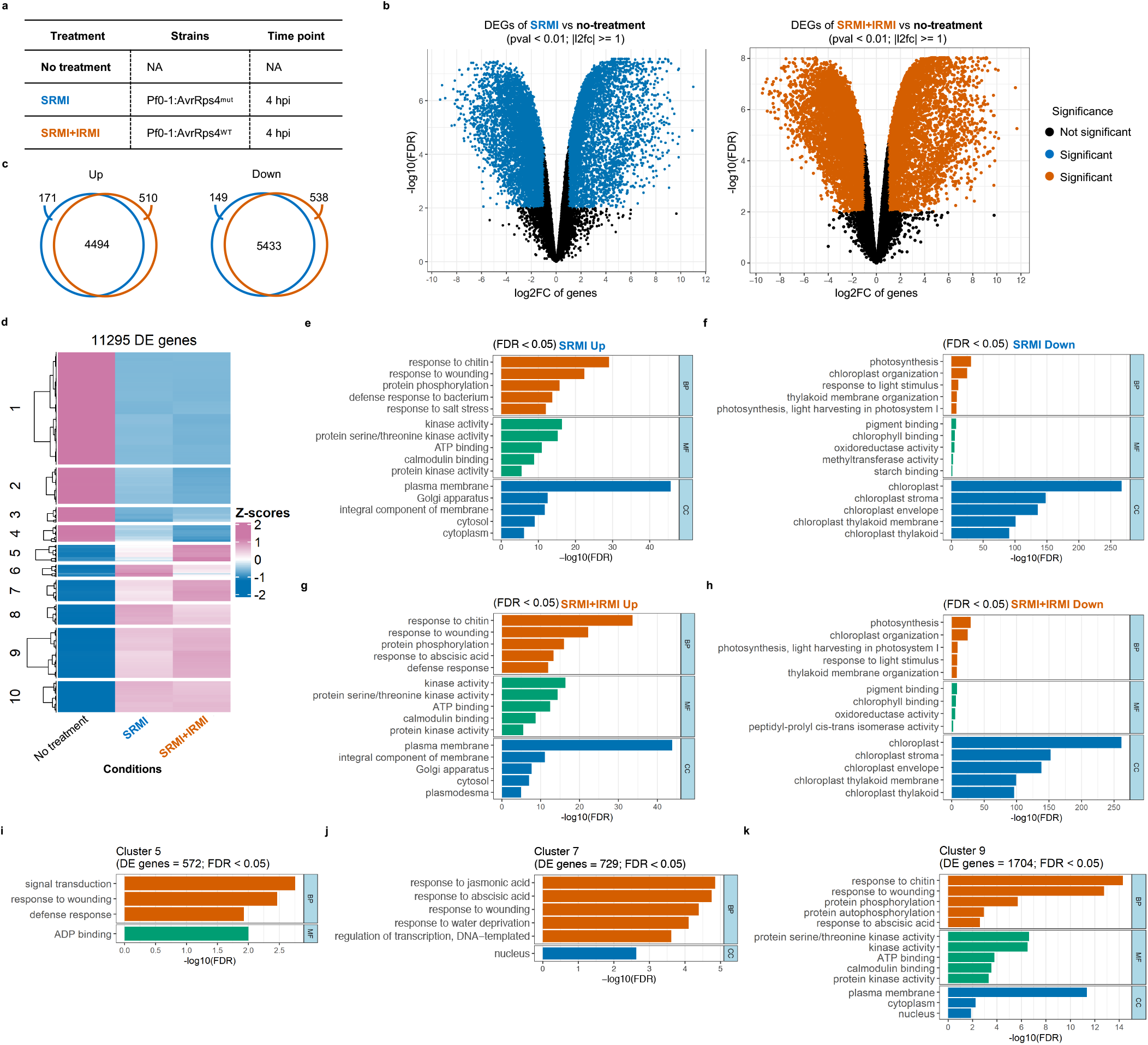
RNA-seq implies differential gene expression during the activation of SRMI and ‘SRMI+IRMI’. **(a)** Simplified table for the information of experimental design for RNA-seq. Three conditions include SRMI, ‘SRMI+IRMI’ with ‘No treatment’ as control. SRMI is implied by the infiltration of Pf0-1 EtHAn carrying mutant AvrRps4 (KRVY 135-138 to AAAA) (Pf0-1:AvrRps4^mut^), which cannot activate NLRs. ‘SRMI+IRMI’ is activated by the infiltration of Pf0-1:AvrRps4^WT^ that is recognized by NLRs RRS1/RPS4 and RRS1B/RPS4B. All sample with infiltration were collected at 4 hours post infiltration (hpi) for RNA extraction and RNA-seq. **(b)** Volcano plots for differentially expressed genes (DEGs) in different conditions. The left panel shows SRMI vs ‘No treatment’ and the right shows ‘SRMI+IRMI’ vs ‘No treatment’. **(c)** Venn diagram illustration DEG comparison between SRMI and ‘SRMI+IRMI’. The left Venn diagram shows comparison of upregulated genes between SRMI and ‘SRMI+IRMI’ and the right shows comparison of downregulated genes. Lists of genes shown in the Venn diagram can be found in Supplementary Table 4. **(a-c)** Treatments are color coded with color-blind friendly palette. ‘No treatment’ in black, SRMI in blue and ‘SRMI+IRMI’ in vermillion. **(d)** Heat map of DEGs based on row z-scores comparing three indicated conditions. Upregulated genes are colored in reddish purple, and downregulated genes are colored in blue. We only set up to 10 clusters. Quantifications of DEGs and related statistics can be found in Supplementary Table 5. **(e-f)** Gene ontology (GO) enrichment for DEGs in SRMI vs ‘No treatment’. **(g-h)** GO enrichment for DEGs in ‘SRMI+IRMI’ vs ‘No treatment’. **(e-h)** Top 5 GO terms were shown in different enrichment analysis, including ‘biological process (BP)’ in vermillion, ‘molecular function (MF)’ in bluish green and ‘cellular compartment (CC)’ in blue (same color code for the rest of the figures in this study). The cut-off for the false discovery rate (FDR) was selected for less than 0.05 (FDR < 0.05) as significant enrichment. The y-axis indicates the value of -log10(FDR) for each GO terms shown. **(i-k)** GO analysis for DEGs in Cluster 5, 7 and 9 indicated the heat map in (d), respectively. Numbers of DEGs in Cluster 5, 7 and 9 are 572, 729 and 1704, respectively. Expression of genes in Cluster 5, 7 and 9 are genes that have higher expression in ‘SRMI+IRMI’ than in SRMI alone, or IRMI-boosted genes (IBGs). FDR < 0.05.

**Supplementary Figure 7.**
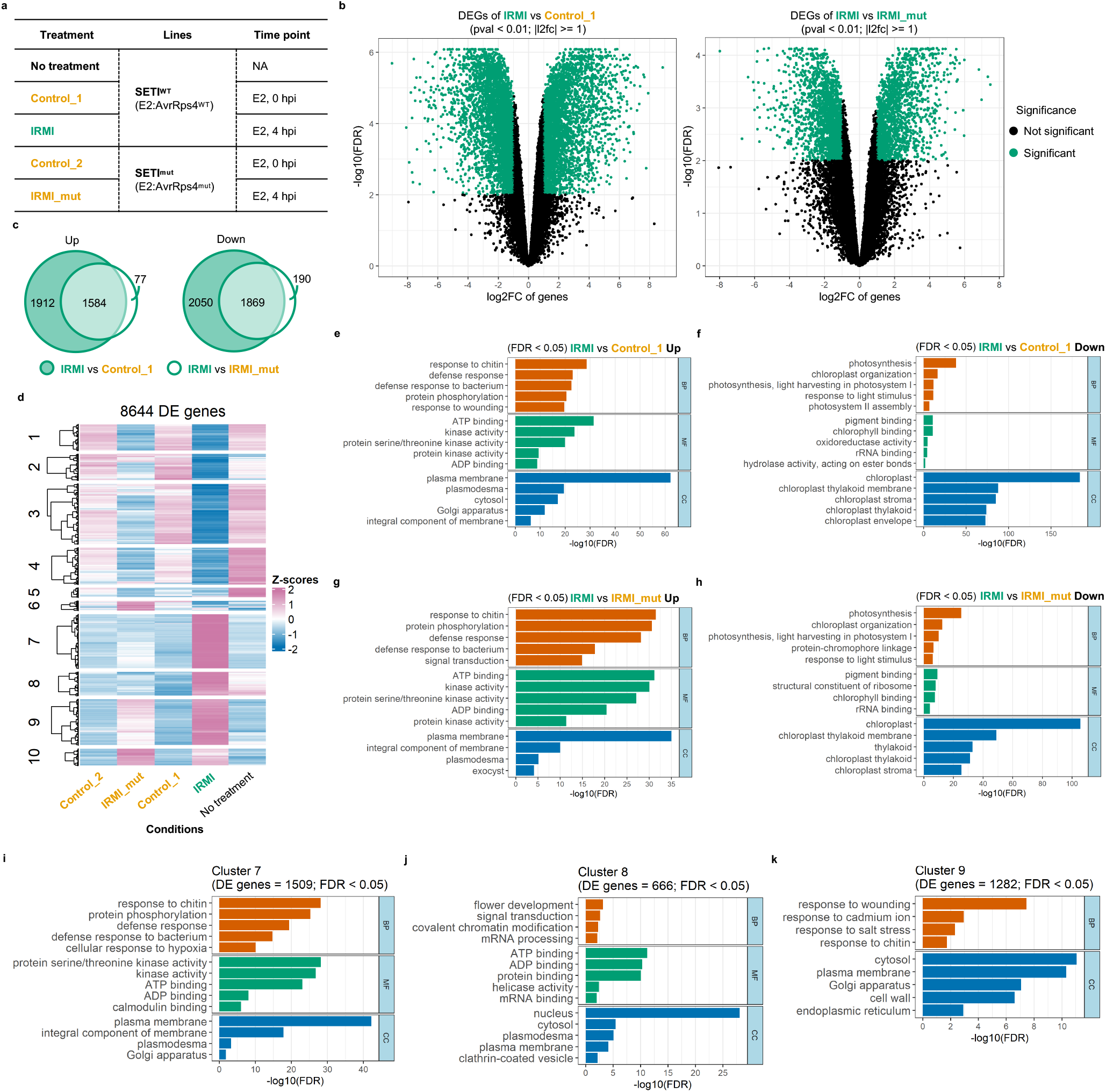
RNA-seq implies differential gene expression during the activation of IRMI. **(a)** Simplified table for the information of experimental design for RNA-seq. IRMI and four different control conditions including ‘No treatment’, Control_1, Control_2 and IRMI_mut are shown here. IRMI is implied by the infiltration of 50 µM β-estradiol (E2) in leaves of SETI^WT^ (inducible AvrRps4 expressing plants, E2:AvrRps4^WT^). ‘No treatment’ in SETI^WT^ serves as a negative control for any concerns regarding possible leakage of the inducible line. Control_1 with E2 infiltrated SETI^WT^ harvest at 0 hpi is for the condition before IRMI activation, so serves as a negative control of IRMI inactive condition. The inducible line SETI^mut^ expressing a mutant AvrRps4 (KRVY135-138AAAA) (E2:AvrRps4^mut^) at 4 hpi of the infiltration of 50 µM E2 serves as a negative control for the concern of side effects of E2 might cause to Arabidopsis plants, and also serve as a negative control for the potential wounding effect introduced by the process of infiltration (IRMI_mut). All sample with infiltration were collected at indicated time points in the table for RNA extraction and RNA-seq. **(b)** Volcano plots for differentially expressed genes (DEGs) in different conditions. The left panel shows IRMI vs ‘Control_1’, which indicates DEGs of before and after IRMI activation. The right panel shows ‘IRMI’ vs ‘IRMI_mut’, which indicates the DEGs of IRMI normalized against wounding. Both contains DEGs that are specific to IRMI. **(c)** Venn diagram illustration DEG comparison between two different comparisons in (b). The left Venn diagram shows comparison of upregulated genes and the right shows comparison of downregulated genes. Lists of genes shown in the Venn diagram can be found in Supplementary Table 6. **(a-c)** Treatments are color coded with color-blind friendly palette. ‘No treatment’ in black. Control_1, Control_2 and IRMI_mut in orange. IRMI in bluish green. **(d)** Heat map of DEGs based on row z-scores comparing three indicated conditions. Upregulated genes are colored in reddish purple, and downregulated genes are colored in blue. We only set up to 10 clusters. Quantifications of DEGs and related statistics can be found in Supplementary Table 7. **(e-f)** GO enrichment for DEGs in IRMI vs Control_1. **(g-h)** GO enrichment for DEGs in IRMI vs IRMI_mut. **(e-h)** Top 5 GO terms were shown in different enrichment analysis, including BP, MF and CC. The cut-off for the false discovery rate (FDR) was selected for less than 0.05 (FDR < 0.05) as significant enrichment. The y-axis indicates the value of -log10(FDR) for each GO terms shown. **(i-k)** GO analysis for DEGs in Cluster 7, 8 and 9 indicated the heat map in (d), respectively. Numbers of DEGs in Cluster 7, 8 and 9 are 1509, 666 and 1282, respectively. Expression of genes in Cluster 7∼9 are genes that have higher expression in IRMI than in all the other control conditions, or IRMI-elevated genes. FDR < 0.05.

**Supplementary Figure 8.**
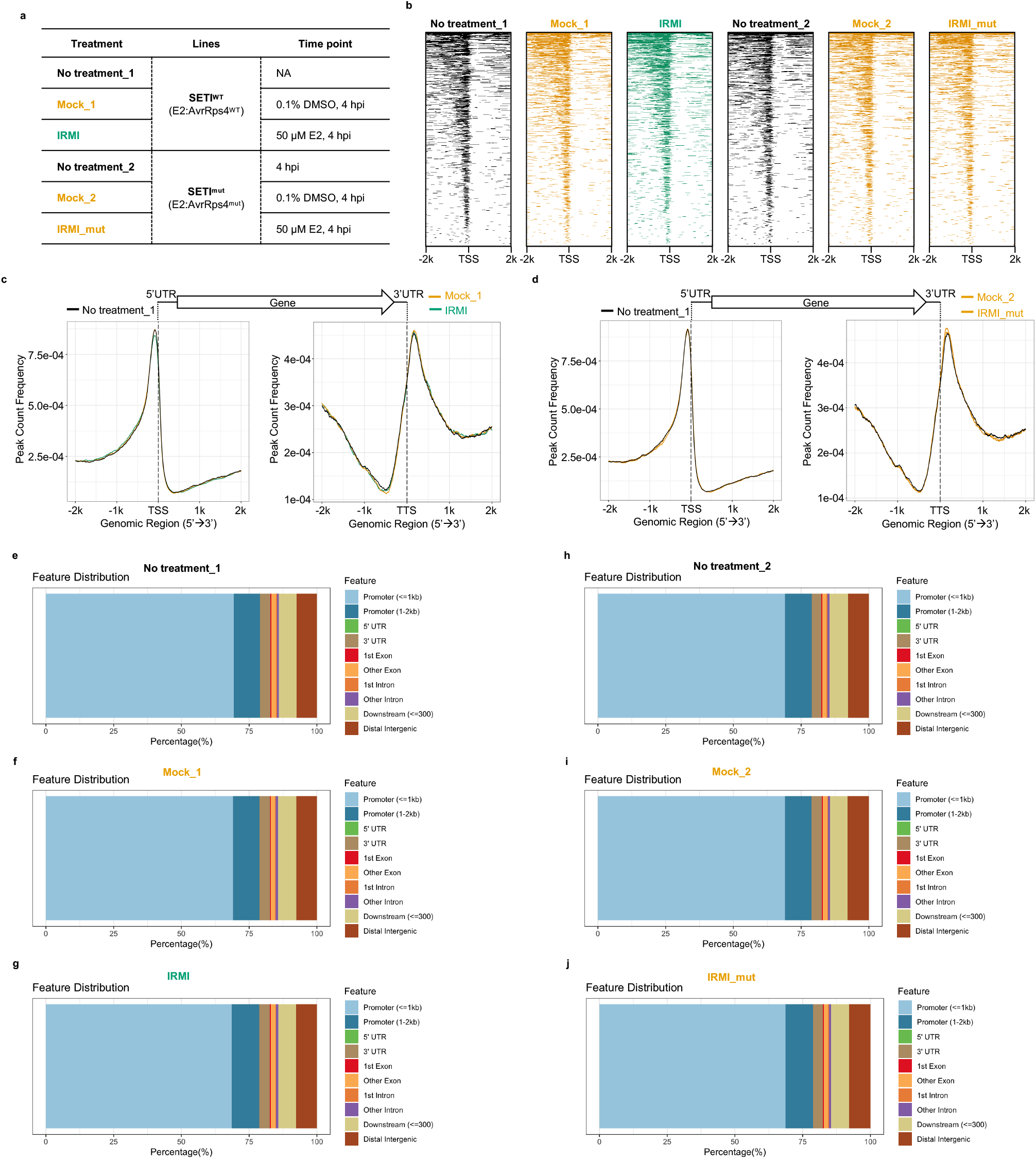
Interrogation of chromatin landscapes activated by IRMI. **(a)** Simplified table for the information of experimental design for FANS-ATAC-seq. IRMI and five different control conditions including ‘No treatment_1’, Mock_1, ‘No treatment_2’, Mock_2 and IRMI_mut are shown here. IRMI is implied by the infiltration of 50 µM β-estradiol (E2) in leaves of SETI^WT^ (inducible AvrRps4 expressing plants, E2:AvrRps4^WT^). ‘No treatment_1’ in SETI^WT^ serves as a negative control for any concerns regarding possible leakage of the inducible line. Mock_1 with 0.1% DMSO infiltrated SETI^WT^ harvest at 4 hpi is a control for wounding caused by infiltration, so serves as a negative control of IRMI inactive condition. The inducible line SETI^mut^ expressing a mutant AvrRps4 (KRVY135-138AAAA) (E2:AvrRps4^mut^) at 4 hpi of the infiltration of 50 µM E2 serves as a negative control for the concern of side effects of E2 might cause to Arabidopsis plants, and also serve as a negative control for the potential wounding effect introduced by the process of infiltration (IRMI_mut). All sample with infiltration were collected at indicated time points in the table and immediately used for FANS-ATAC-seq. **(b)** Heatmaps showing the distribution of accessible regions around the TSS identified by FANS-ATAC-seq (average value of two biological replicates) under four different conditions. Accessible regions are mapped to 2, 000 bp upstream (−2k) or downstream (2k) of TSS as the center. ‘No treatment_1’ and ‘No treatment_2’ are indicated in black; Mock_1, Mock_2 and IRMI_mut are indicated in orange; IRMI is indicated in blueish green. (same color codes apply to the same set of treatments in the rest of this study). **(c)** Distribution of accessible regions around the TSS (left panel) and TTS (right panel) identified from FANS-ATAC-seq with the mean peak counts from two biological replicates indicated in (b). The center of accessible regions was used to produce the distribution plots. **(e-j)** Accessible chromatin regions from all conditions are mapped to different genomic features as indicated.

**Supplementary Figure 9.**
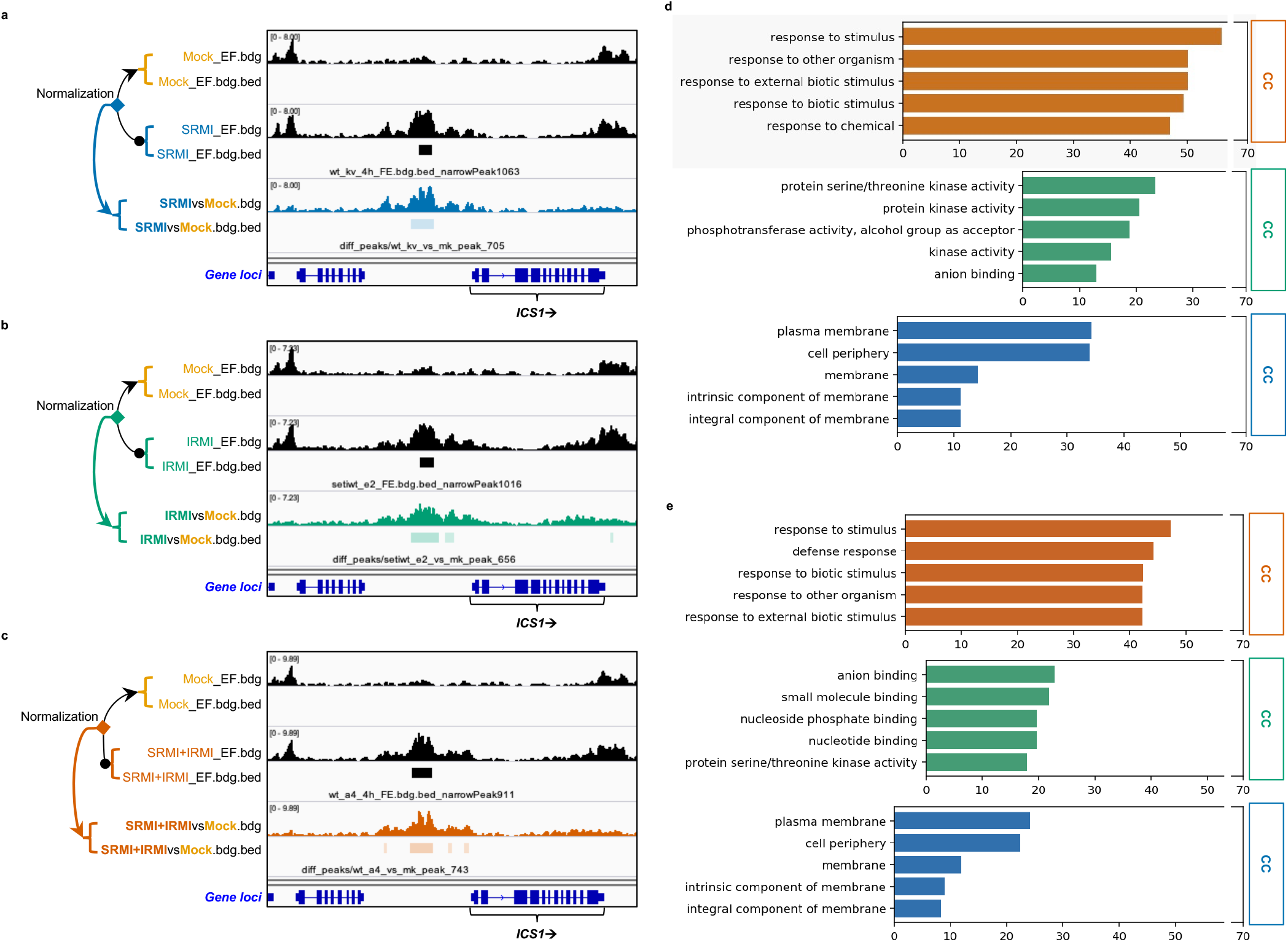
Normalization of accessible chromatin regions for DARs and related GO enrichment. **(a-c)** Schematic demonstration of normalization of accessible chromatin regions for DARs. (a) SRMI, (b) IRMI and (c) ‘SRMI+IRMI’ were normalized to corresponding Mock, respectively. More details can be found in Methods. **(d)** GO enrichment for top 5 listed terms for 1413 genes listed in the intersection of ‘DAR ∩ DEG’ in SRMI and ‘DAR ∩ DEG’ in ‘SRMI+IRMI’ in Fig. 4c. **(e)** GO enrichment for top 5 listed terms for 947 genes listed in the intersection of DARs and DEGs in IRMI in Fig. 4d.

**Supplementary Figure 10.**
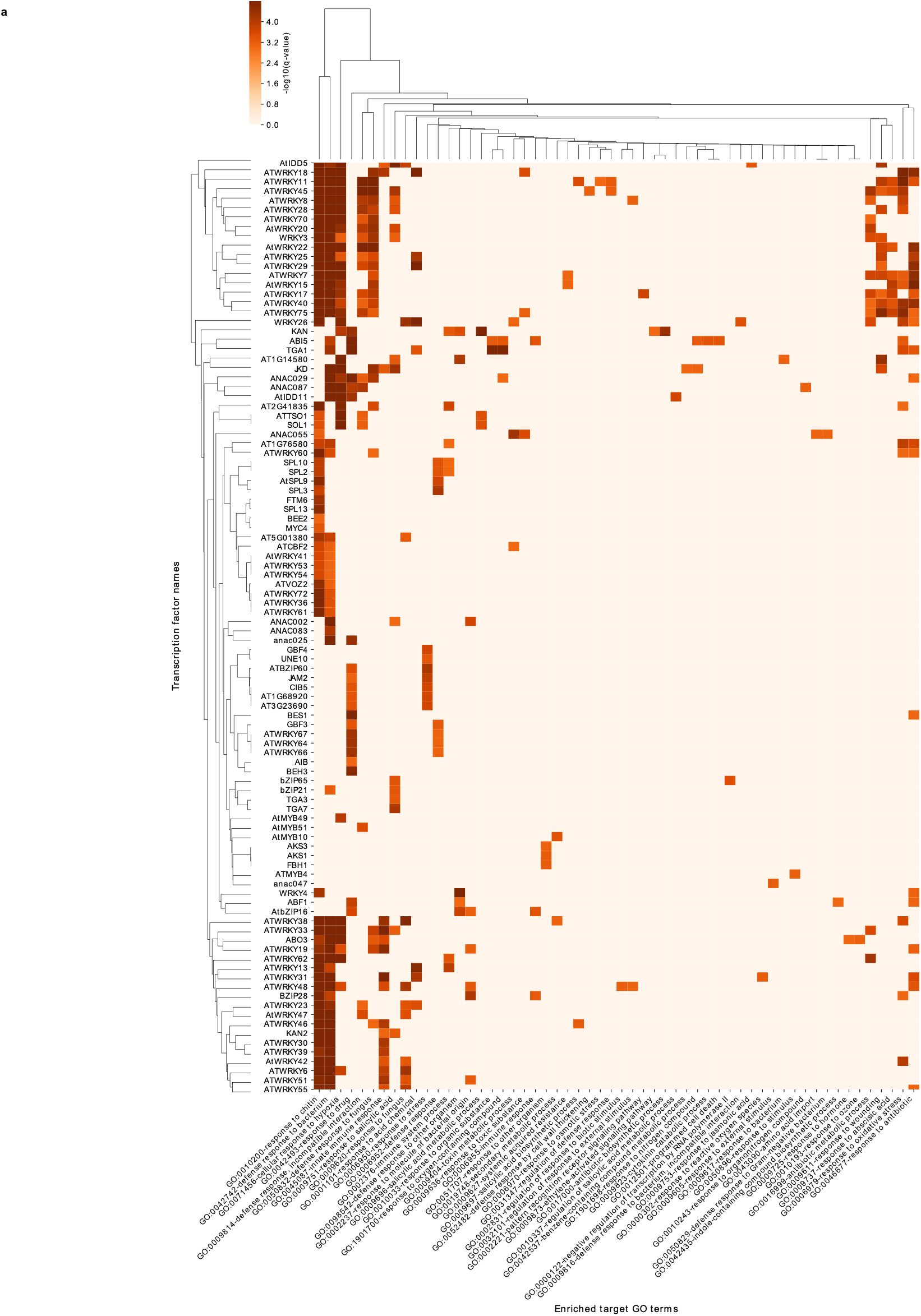
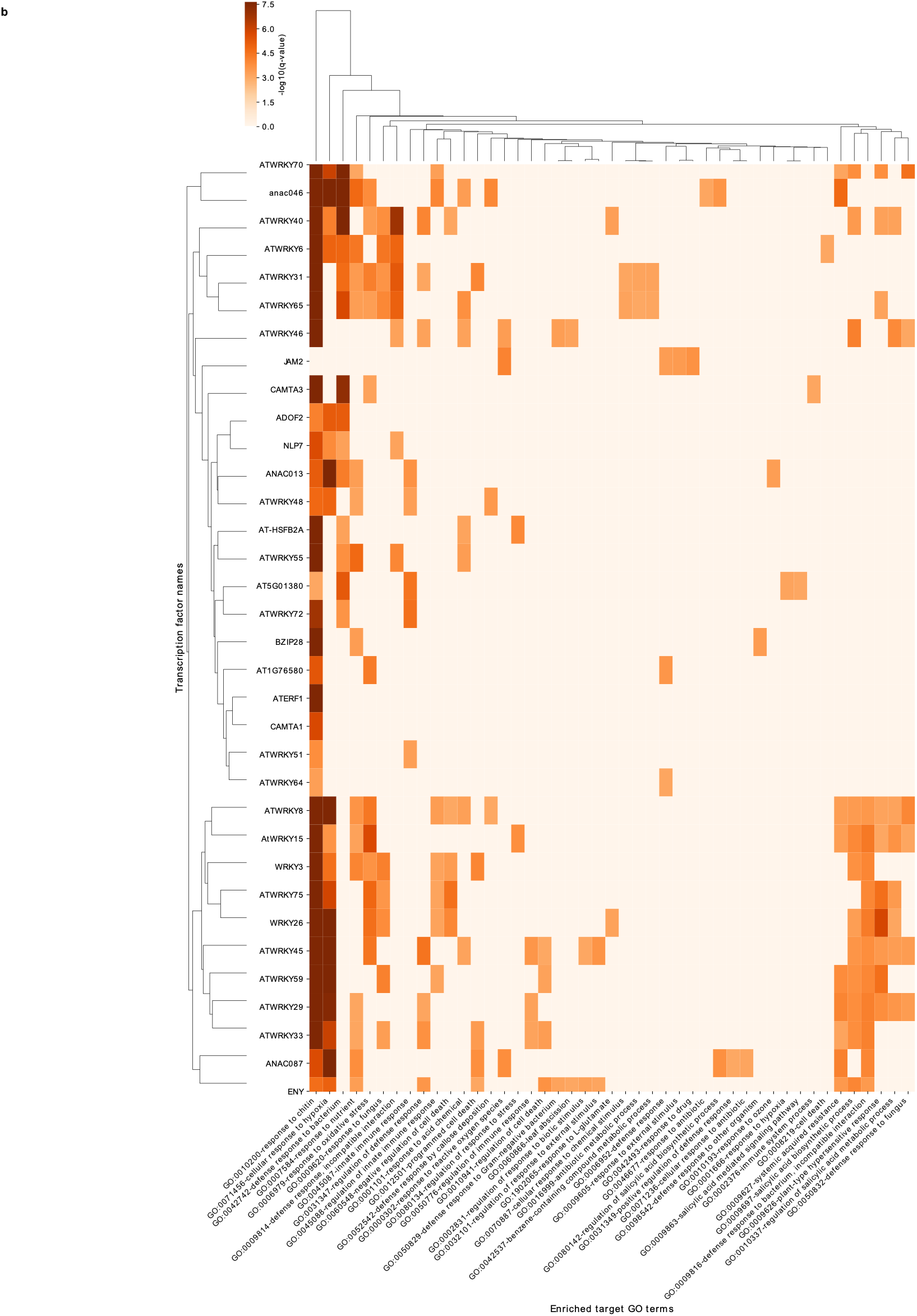
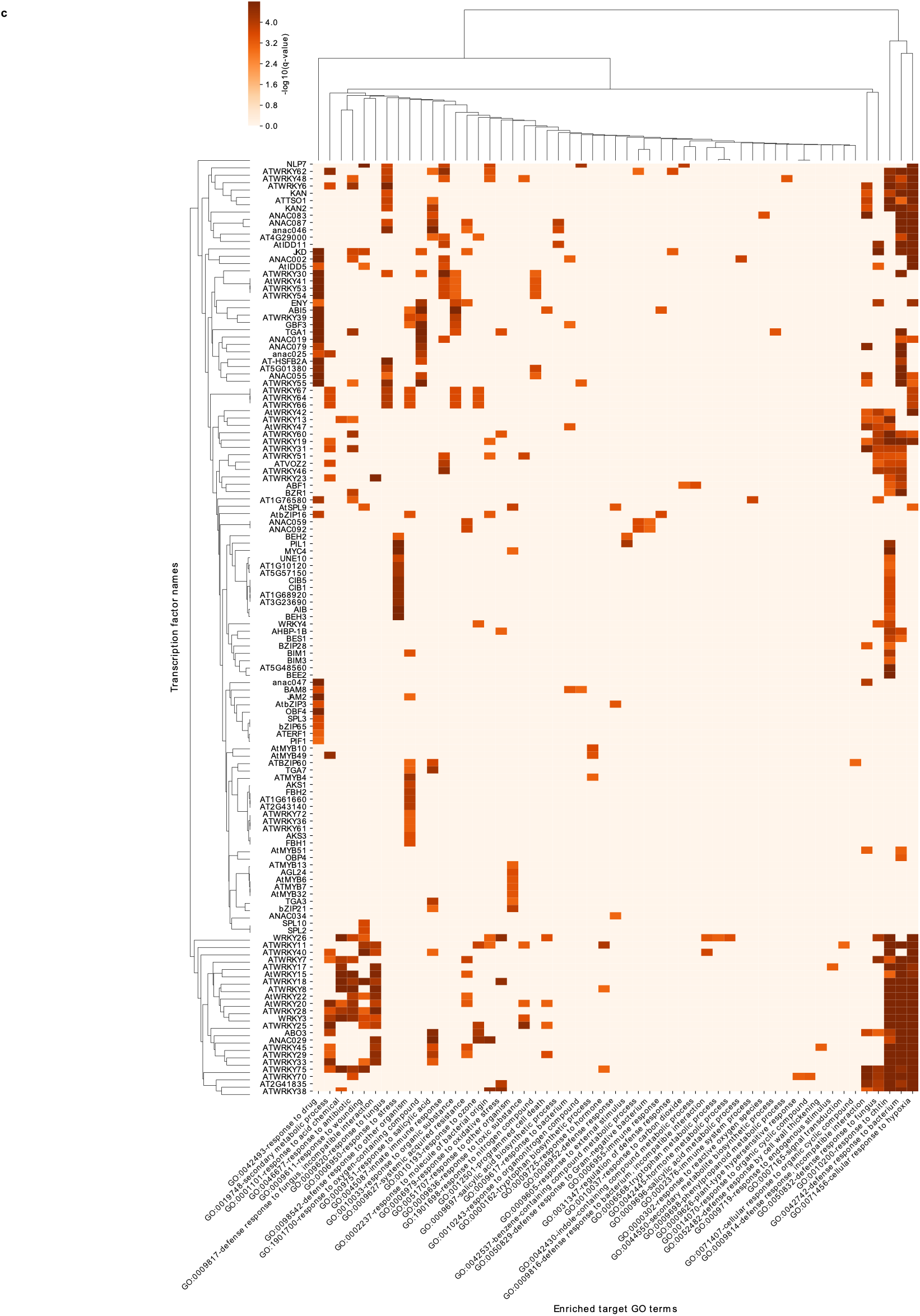
TF target gene GO enrichment heatmaps. **(a - c)** Heatmaps showing the GO enrichment -log10(q-value) of the putative target genes for each TF. **(a)** SRMI. **(b)** IRMI, **(c)** ‘SMRI+IRMI’.

**Supplementary Figure 11.**
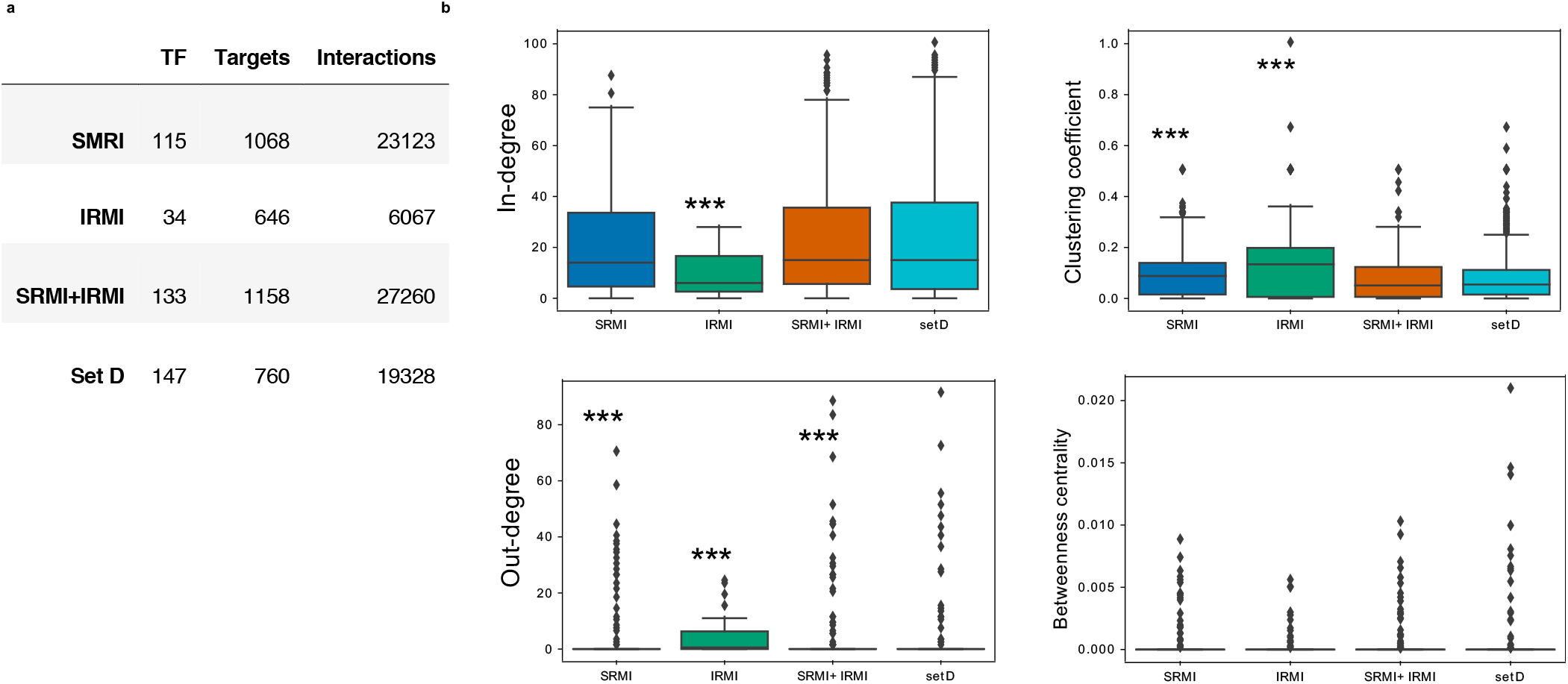
Network centrality parameters. **(a)** Number of TFs in each network, the number of putative target genes they control and the interactions that there are between them for each condition and for set D. Set D is a control network that contains the DE TFs shared in the three conditions and all the non-redundant target genes and interactions that each of these TFs have in the three networks. **(b)** In-degree, out-degree, clustering coefficient and betweenness centrality computations for all the nodes in each network. The statistical significance was determined pairwise against the set D network with a two-sided Mann-Whitney U test (*** means p-value < 0.001). The network parameters were computed using NetworkX python package v2.4.

**Supplementary Figure 12.**
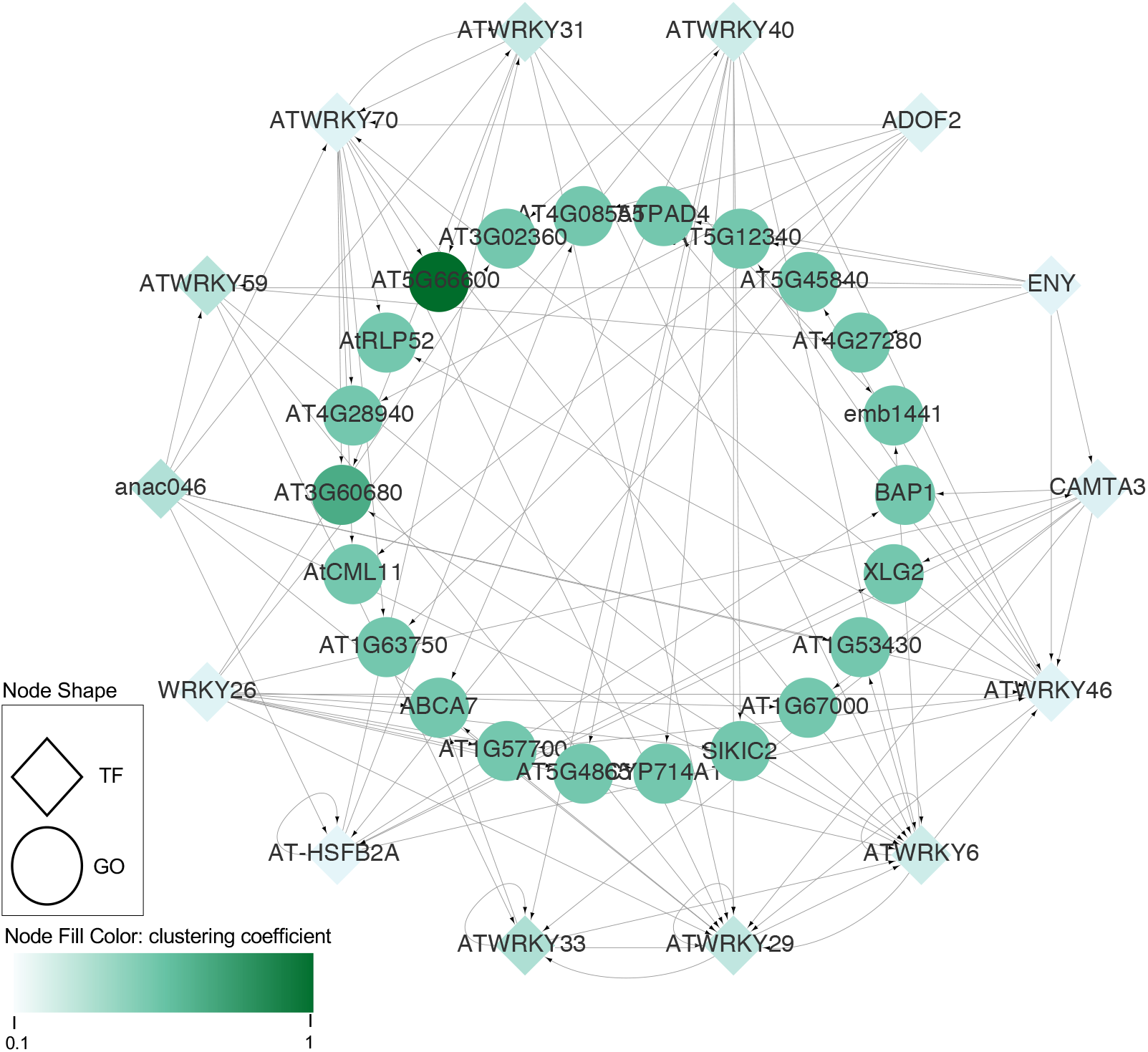
IRMI network for targets with high clustering coefficient. The nodes from the IRMI network that have a clustering coefficient ≥ 0.5 and the incoming TFs that putatively control their expression are represented in the network. The nodes are colored according to their clustering coefficient.

## Supplementary Tables

**Supplemenatry Table 1: Dual index primers for multiplexing; associated with Supplementary Fig. 2 and Fig. 3**

**Supplementary Table 2: Destribution of enriched accessible chromatins on different genomic features under different treatments in Supplementary Fig.5**

Supplementary Table S2.1: Destribution of enriched accessible chromatins on different genomic features in ‘No treatment’

Supplementary Table S2.2: Destribution of enriched accessible chromatins on different genomic features in Mock

Supplementary Table S2.3: Destribution of enriched accessible chromatins on different genomic features in SRMI

Supplementary Table S2.4: Destribution of enriched accessible chromatins on different genomic features in ‘SRMI+IRMI’

**Supplementary Table 3: Accessible chromatin regions enriched in different treatment groups and intersections between groups in Fig.1c**

**Supplementary Table 4: Statistics of differentially expressed genes listed in Supplementary Fig. 6c**

Supplementary Table S4.1: Statistics of up-regulated genes in Supplementary Fig. 6c

Supplementary Table S4.2: Statistics of down-regulated genes in Supplementary Fig. 6c

**Supplementary Table 5: Expression value and cluster information of genes listed in Supplementary Fig. 6d**

Supplementary Table S5.1: Transcripts Per Million (TPM) for gene expression profiling related to Supplementary Fig. 6d

Supplementary Table S5.2: Gene targets enriched in each cluster listed in Supplementary Fig. 6d

**Supplementary Table 6: Statistics of differentially expressed genes listed in Supplementary Fig. 7c**

**Supplementary Table 7: Expression value and cluster information of genes listed in Supplementary Fig. 7d**

Supplementary Table S7.1: Transcripts Per Million (TPM) for gene expression profiling related to Supplementary Fig. 7d

Supplementary Table S7.2: Gene targets enriched in each cluster listed in Supplementary Fig. 7d

**Supplementary Table 8: Destribution of enriched accessible chromatins on different genomic features under different treatments in Supplementary Fig. 8**

Supplementary Table S8.1: Destribution of enriched accessible chromatins on different genomic features in ‘No treatment_1’

Supplementary Table S8.2: Destribution of enriched accessible chromatins on different genomic features in Mock_1

Supplementary Table S8.3: Destribution of enriched accessible chromatins on different genomic features in IRMI

Supplementary Table S8.4: Destribution of enriched accessible chromatins on different genomic features in ‘No treatment_2’

Supplementary Table S8.5: Destribution of enriched accessible chromatins on different genomic features in Mock_2

Supplementary Table S8.6: Destribution of enriched accessible chromatins on different genomic features in IRMI_mut

**Supplementary Table 9: Gene lists and their GO terms enriched in DARs and DEGs data integration; related to Fig. 4 and Supplementary Fig. 9**

Supplementary Table 9.1: DARs and DEGs in different contrast groups and intersections between groups in Fig. 4

Supplementary Table 9.2: Enriched GO terms associated with ‘DAR ∩ DEG’ shared by SRMI and ‘SRMI+IRMI’

Supplementary Table 9.3: Enriched GO terms associated with ‘DAR ∩ DEG’ in ‘IRMI vs Mock’

**Supplementary Table 10: Information for gene regulatory networks learned from ATAC-seq and RNA-seq under different immune conditions; related to Fig. 5, 6 and Supplementary Fig. 10-12**

Supplementary Table 10.1: Transcription factors and their targets inferred from ATAC-seq and transcription factor binding motif information

Supplementary Table 10.2: Gene regulatory network matrix linking transcription factors to their targets

